# TRPM2-CaMKII signaling drives excessive GABAergic synaptic inhibition following ischemia

**DOI:** 10.1101/2023.09.07.556550

**Authors:** Amelia M. Burch, Joshua D. Garcia, Heather O’Leary, Ami Haas, James E. Orfila, Erika Tiemeier, Nicholas Chalmers, Katharine R. Smith, Nidia Quillinan, Paco S. Herson

**Affiliations:** Department of Anesthesiology, Neuronal Injury & Plasticity Program, University of Colorado School of Medicine, Anschutz Medical Campus, 12801 E. 17^th^ Ave., Aurora CO 80045, USA; Department of Pharmacology, University of Colorado School of Medicine, Anschutz Medical Campus, 12800 E. 19^th^ Ave., Aurora CO 80045, USA; Department of Neurological Surgery, The Ohio State University College of Medicine, 460 W. 12^th^ Ave., Columbus OH 43210, USA

**Keywords:** GABA(A) receptor, cerebral ischemia, inhibitory synapse, TRPM2, CaMKII, hippocampus, LTP, E/I balance, gephyrin

## Abstract

Following an ischemic insult to the brain, there is an acute loss of GABAergic inhibitory synapses and an increase in excitatory/ inhibitory (E/I) imbalance that drives neuronal hyperexcitability. It is unknown whether this E/I imbalance persists at delayed timepoints and contributes to chronic impairments in memory and long-term potentiation (LTP) in the hippocampus following ischemic brain injury. Here, we reveal a shift to reduced E/I ratio in hippocampal CA1 neurons via a persistent increase in postsynaptic GABA_A_ receptor mediated inhibitory responses and clustering days after a global ischemic insult. This enhancement of postsynaptic inhibitory function and clustering required activation of the Ca^2+^-permeable TRPM2 ion channel and the Ca^2+^-dependent kinase, CaMKII. Thus, we propose a mechanism in which acute downregulation of GABA_A_ receptors is followed by a strengthening of inhibitory synapses at delayed periods after ischemia. Targeting this mechanism has therapeutic potential to recover hippocampal plasticity and cognitive function post-ischemia.

**GRAPHICAL ABSTRACT:** 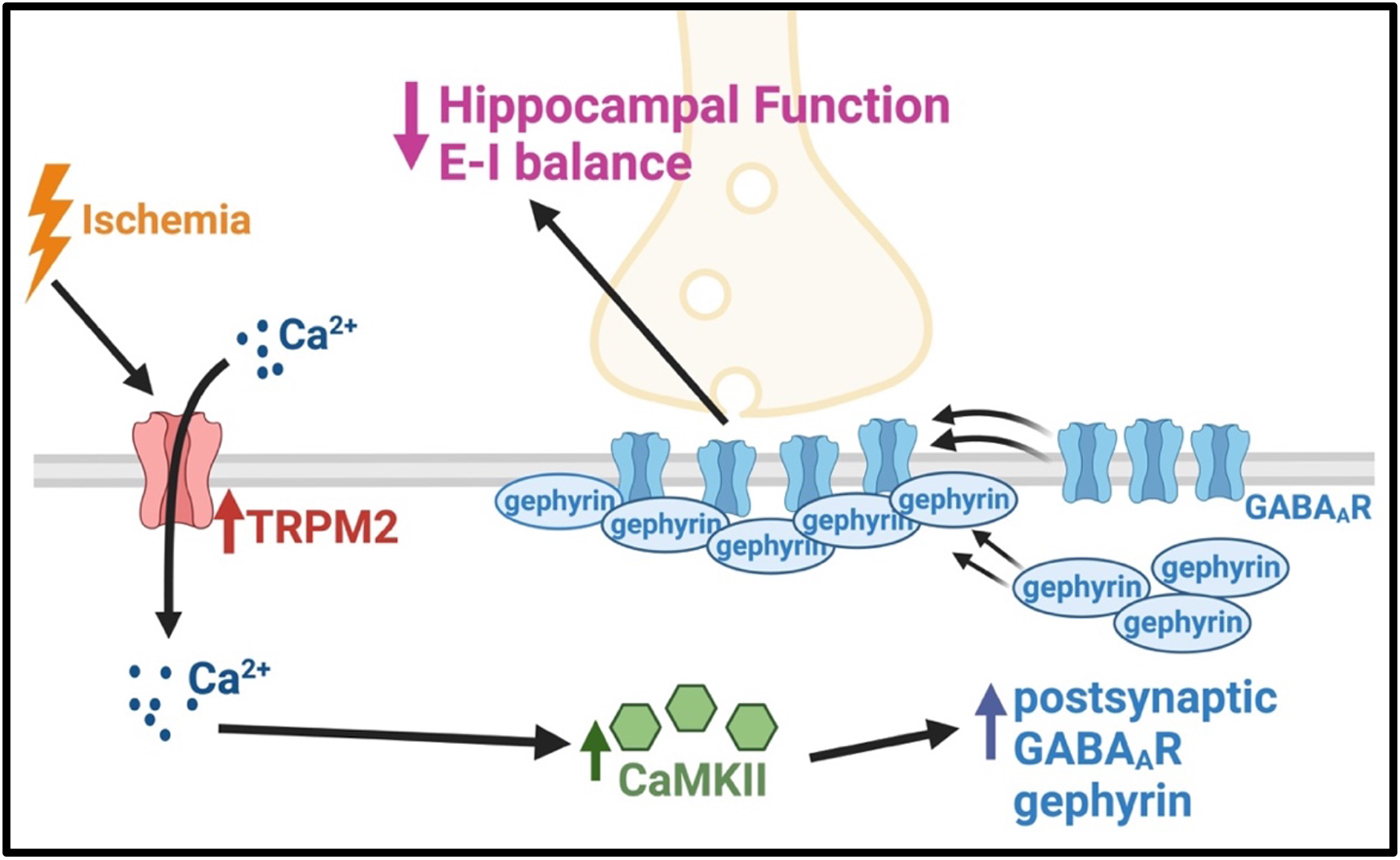

## INTRODUCTION

Hippocampal dysfunction occurs as a consequence of brain injury, leading to impairments in learning and memory. Notably, cerebral ischemia has profound effects on hippocampal function due to the high metabolic requirements of the hippocampus (Neumann et al., 2013; Petito et al., 1987; Schmidt-Kastner and Freund, 1991). Ischemic insults immediately trigger a cascade of signaling events which culminate in neuronal cell death. Specifically, excitotoxicity is one pathway, ubiquitous to numerous neurological disorders, which drives neuronal demise via excessive glutamate release and resultant activation of AMPA- and NMDA-type glutamate receptors. Thus, considerable research efforts have concentrated on mitigating these acute excitotoxic effects through NMDA and/ or AMPA receptor blockade (Kostandy, 2012; Turski et al., 1998; Walters et al., 2005; Wu and Tymianski, 2018). While this neuroprotective approach effectively reduces neuronal cell death in animal models, clinical trials have shown limited improvement in long-term functional recovery in patients likely due to the narrow therapeutic window required for these therapies (Cheng et al., 2004; Katz et al., 2022; Wahlgren and Ahmed, 2004).

Neurorestoration is an alternative strategy that aims to restore neural circuits perturbed by neural injury, by enhancing synaptic function and plasticity in surviving neurons, allowing for treatment within a broader therapeutic timeframe (Azad et al., 2016; Escobar et al., 2019). Indeed, this strategy can be reliably tested in a murine model of global cerebral ischemia (GCI), cardiac arrest/ cardiopulmonary resuscitation (CA/CPR). Despite completion of cell death processes within days of the initial insult, we observed sustained impairments of long-term potentiation (LTP) in surviving neurons within the hippocampus, correlating with learning and memory deficits post-CA/CPR (Dietz et al., 2019; Orfila et al., 2014). More recently, we demonstrated that delayed inhibition of the Ca^2+^-permeable, transient receptor potential melastatin-2 (TRPM2) ion channel reverses impairments in hippocampal LTP and hippocampal-dependent behavioral tasks (Dietz et al., 2019; Dietz et al., 2021), implicating TRPM2 as a potential target for neurorestorative therapy following ischemia. However, the precise molecular mechanisms underlying hippocampal dysfunction and the contribution of ongoing TRPM2 activity to LTP impairment remain elusive.

Extensive studies have emphasized the importance of maintaining a balance between excitatory and inhibitory (E/I) signaling for optimal neuronal function and induction of LTP mechanisms (Smith and Kittler, 2010; Vogels et al., 2011). One possible mechanism of ischemia-induced LTP impairment is disrupted E/I balance through reduced excitatory glutamatergic transmission mediated by AMPA- or NMDA-type glutamate receptors. However, we found that both the function and expression of these receptors were mostly unaltered 7 days following CA/CPR (Orfila et al., 2018; Orfila et al., 2014), suggesting changes in excitatory signaling likely do not account for the ongoing LTP deficits. Thus, it is plausible that enhanced inhibitory GABAergic function may reduce the E/I ratio and contribute to LTP impairments. Indeed, prior work indicates that inhibitory signaling directly impacts excitatory LTP (Leao et al., 2012; Steele and Mauk, 1999; Udakis et al., 2020; Williams and Holtmaat, 2019). Specifically, GABAergic synaptic inhibition plays a critical role in E/I balance by regulating circuit and neuronal excitability (Chiu et al., 2019). A mechanism by which strengthening of GABAergic synapses is achieved is through increased clustering of GABA_A_ receptors (GABA_A_Rs) at synaptic sites (Luscher et al., 2011; Nusser et al., 1997). Disruptions in the trafficking of GABA_A_Rs to the synapse have deleterious effects on excitatory synaptic plasticity and are implicated in various neurologic conditions (Mele et al., 2019; Smith and Kittler, 2010). In the context of cerebral ischemia, there is accumulating evidence that excessive activation of synaptic (phasic) (Hiu et al., 2016) and extrasynaptic (tonic) GABA_A_Rs are deleterious to functional recovery in the post-acute phase, days following the completion of cell death processes (Carmichael, 2012; Clarkson et al., 2010; Orfila et al., 2019). However, the precise upstream mechanisms driving this enhancement and whether these pathways contribute to sustained hippocampal synaptic plasticity and cognitive deficits remain poorly defined. In light of the established role of GABAergic inhibition in modulating excitatory synaptic plasticity (Leao et al., 2012; Udakis et al., 2020; Williams and Holtmaat, 2019), we tested the effect of ischemia on synaptic GABAergic inhibition and whether TRPM2 reverses LTP impairments via modulation of GABA_A_R function.

Here, we present data demonstrating a shift in the balance of E/I signaling in the hippocampus after CA/CPR. At acute timepoints, we found a decrease in GABA_A_R clustering. However, in the post-acute phase following the completion of cell death processes, we observed a sustained increase in postsynaptic inhibition, leading to a reduction in E/I balance. Using a combination of *in vitro* and *in vivo* approaches, we rigorously identified the TRPM2 ion channel as a mediator of augmented inhibitory function. We also provide compelling evidence that TRPM2 and Ca^2+^/calmodulin-dependent protein kinase II (CAMKII) activation are necessary for the sustained enhancement of postsynaptic GABAergic inhibition, highlighting potential therapeutic targets to improve functional recovery following brain ischemia.

## MATERIALS & METHODS

### Key Resources Table

| REAGENT or RESOURCE | SOURCE | IDENTIFIER |
| --- | --- | --- |
| **Antibodies** |  |  |
| GABA<sub>A</sub>R-γ2 (imaging) | Synaptic Systems | Cat. #224 004; AB_10594245 |
| Gephyrin (3B11; imaging) | Synaptic Systems | Cat. #147 111; AB_2619837 |
| VGAT (rabbit; imaging) | Synaptic Systems | Cat. #131 003; AB_887869 |
| Anti-guinea pig Alexa Fluor - 568 | Life Technologies | Cat. #11075; AB_141954 |
| Anti-mouse Alexa Fluor - 488 | Life Technologies | Cat. #21202; AB_141607 |
| Anti-rabbit Alexa Fluor - 647 | Life Technologies | Cat. #31573; AB_2536183 |
| Gephyrin (WB) | Synaptic Systems | Cat. #147 111; AB_2619837 |
| phosphoCaMKII T286 | PhosphoSolutions | Cat. #p1005-286; AB_2492051 |
| αCaMKII | Gift from KU Bayer | |
| HRP- conjugated β-Actin | Sigma-Aldrich | Cat. #3854 |
| HRP-conjugated goat anti-mouse | Jackson ImmunoResearch | Cat. #115-035-174; AB_2338512 |
| HRP-conjugated goat anti-rabbit | ThermoScientific | Cat. #31460; AB_228341 |
| **Chemicals, Reagents and Kits** |  |  |
| Picrotoxin | Tocris | Cat. #1128 |
| Qx314 | Tocris | Cat. #1014 |
| DNQX | Tocris | Cat. #0189 |
| Clotrimazole | Sigma-Aldrich | Cat. #6019 |
| **Experimental Models: Organisms/Strains** |  |  |
| Rat, Sprague Dawley Charles River | Charles River | RRID: RGD_734476 |
| Mice, C57BL/6 | Jackson Laboratory |  |
| **Software** |  |  |
| Prism 9 | GraphPad | https://www.graphpad.com/scientific-software/prism/ |
| ImageJ | NIH | https://imagej.nih.gov/ij/ |

### Experimental Model and Subject Details

#### Animals

All studies conformed to the requirements of the National Institutes of Health *Guide for the Care and Use of Laboratory Animals* and were approved by the Institutional Animal Care and Use subcommittee of the University of Colorado, Denver AMC. C57BL/6 mice were bred in house in the Animal Resource Center at the University of Colorado Anschutz Medical Campus and monitored regularly for health. Mice were weaned between postnatal day 21 and 28 (P21-28) and housed in micro-isolator cages on a 14:10 light: dark cycle with water and chow available *ad libitum*.

#### Cardiac arrest/ cardiopulmonary resuscitation

Male mice that were approximately 8-12wks old were subjected to either cardiac arrest and cardio-pulmonary resuscitation (CA/CPR) or sham procedures as described previously (Deng et al., 2017; Dietz et al., 2019). Briefly, mice were anesthetized with 3% isoflurane. Mice were intubated and connected to a mouse ventilator set to 160 breaths per min. Cardiac function was monitored via electrocardiography, and pericranial temperature was maintained at 37.5°C ± 0.2°C using a water-filled coil. Asystolic cardiac arrest was induced by KCl injection via jugular catheter. CPR was begun 6min after induction of cardiac arrest, by slow injection of 0.5–1.0 mL of epinephrine (16μg epinephrine/mL, 0.9% saline), chest compressions at a rate of ∼300min^−1^, and ventilation with 100% oxygen. If return of spontaneous circulation could not be achieved within 3min of CPR, resuscitation was terminated and the mouse was excluded from the study. After surgical procedures, mice were housed individually in micro-isolator cages on heating pads. Post-surgical care included daily saline injections (1 mL) and moist chow for 72hrs. Investigators performed all experiments blind to surgical procedure of the animal, with separate investigator generating the code.

#### Dissociated hippocampal cultures

Primary hippocampal cultures were prepared as described previously(Crosby et al., 2019; Garcia et al., 2021; Rajgor et al., 2020). Briefly, the hippocampi from neonatal rat pups (P0-P1) were dissected and dissociated in papain. The isolated neurons were then seeded in MEM supplemented with 10% FBS and penicillin/streptomycin. Cells were plated at a density of 150,000-200,000 cells per 18mm, #1.5 glass coverslip coated with poly-D-lysine. The MEM was replaced with Neurobasal (NB) media (GIBCO) supplemented with B27 (GIBCO) and 2mM Glutamax 24hrs after plating. The media were refreshed every 5 days be removing half of the existing media and replacing it with fresh NB media. To restrict the growth of actively dividing cells, mitotic inhibitors (uridine fluoro deoxyuridine) were introduced on day 5. The cultures were maintained at 37°C with 5% CO_2_ for a period of 13-14 days before conducting OGD experiments.

### Method Details

#### Hippocampal slice preparation

Hippocampal slices were prepared 7 days post-surgical procedures. Mice were anesthetized with 3% isoflurane in an oxygen enriched chamber then trans-cardially perfused with oxygenated ice-cold artificial Cerebral Spinal Fluid (aCSF) containing in mM: 126 NaCl, 25 NaHCO_3_, 12 glucose, 2.5 KCl, 2.4 CaCl_2_, 1.3 NaHPO_4_, and 1.2 MgCl_2_. Horizontal slices (300 µM) were cut in aCSF supplemented with 9mM MgSO_4_ and continuous oxygenation using a Vibratome 1200 (Leica) and transferred to a holding chamber containing aCSF warmed to 33°C. After 30min, slices recovered for an additional 30min at room temperature (RT), prior to electrophysiology recordings.

#### Whole cell patch clamp electrophysiology

CA1 pyramidal neurons were visualized using infrared digital interference contrast optics under 63x magnification and patch-clamped in whole cell configuration. Borosilicate glass recording electrodes were pulled with either a Narishige or Sutter P-97 Flaming/Brown electrode puller (Sutter Instruments) with a resistance of 2.5-4.0MΩ. Recording solution was exchanged at a flow rate of approximately 2mL per min at RT. Responses were amplified and filtered at 5 kHz (Multiclamp 700B) and digitized at 20 kHz (Digidata 1440A and Clampex 10.7). No series resistance compensation nor junction potential corrections were performed. Access resistance was monitored by delivering a -5mV voltage step, and any experiments in which access resistance was above 30MΩ or changed more than 20% between the onset and completion of the experiment were not used for experimental analyses.

Excitatory/inhibitory balance experiments were conducted by recording excitatory postsynaptic currents (EPSCs) at -70mV and inhibitory postsynaptic currents (IPSCs) at 0mV from the same cell with an internal solution containing (in mM) 135 CsMeSO_4_, 10 HEPES, 10 BAPTA, 5 Qx314, 4 Na_2_ATP, 4 MgCl_2_, 0.3 NaGTP, and pH 7.25 with 1M CsOH. Responses were evoked using a monopolar electrode in the stratum radiatum to stimulate Schaffer collateral-commissurals (SCC) apical dendrites using a constant current source (Digitimer Ltd). Episodic mode was used to record EPSCs and IPSCs responses evoked every 20s for a minimum of 2min each. Spontaneous inhibitory postsynaptic currents (sIPSCs) were recorded at -60mV in gap free mode, with an internal solution containing (in mM) 140 CsCl, 10 NaCl, 10 HEPES, 5 EGTA, 0.5 CaCl_2_, 2 MgATP, and 5 Qx314 in aSCF containing 10µM DNQX to block AMPA currents(Banks et al., 1998). The EGTA and CaCl_2_ in the internal solution was replaced with 10mM BAPTA to examine the role of Ca^2+^ signaling in postsynaptic neuron sIPSCs following CA/CPR. To determine the role of CaMKII activity in postsynaptic neurons sIPSCs following CA/CPR, the internal solution was supplemented with 5µM tatCN19o(Barcomb et al., 2015) (gift from KU Bayer). To assess the role of TRPM2 activation in inhibitory function, 2µM tatM2NX(Cruz-Torres et al., 2020; Dietz et al., 2019) was bath applied for at least 20-min prior to and during recording. For clotrimazole experiments, baseline was recorded for 3min, and clotrimazole (20µM)(Verma et al., 2012) was perfused onto the slice for 12min. Access resistance was monitored every 3min. Recordings which exhibited >20% drift in access resistance were discarded. The final 3min of clotrimazole recordings were used for statistical comparison to baseline.

#### Field electrophysiology

Hippocampal slices were placed in a heat-controlled interface chamber perfused with aCSF at a rate of 1.5mL/min at 32°C. Responses were evoked using an insulated tungsten bipolar stimulating electrode placed in stratum radiatum to stimulate Schaffer collateral-commissurals (SCC), and recorded with a glass electrode containing 150mM NaCl placed in the distal dendrites of CA1 pyramidal cell layer. Analog fEPSPs were amplified (1000×) and filtered through a preamplifier (Grass Model P511) 0.03 Hz to 1.0 kHz, digitized at 10 kHz and stored on a computer for later off-line analysis (Datawave Technologies). The derivative (dV/dT) of the initial fEPSP slope was measured. The fEPSPs were adjusted to 50% of the maximum slope and test pulses were evoked every 20s. Paired pulse responses were recorded using a 50-ms interpulse interval (20 Hz) and expressed as a ratio of the slopes of the second pulse over the first pulse. Picrotoxin (5μM)(Costa and Grybko, 2005) was applied for at least 20-min during acquisition of a stable fEPSP baseline. Following the baseline recording, we washed out the picrotoxin with normal aCSF and delivered theta burst stimulation (TBS), which included a train of four pulses delivered at 100 Hz in 30-ms bursts repeated 10 times with 200-ms interburst intervals. Following TBS, the fEPSP was recorded for 60min. The averaged 10-min slope from 50 to 60min after TBS was divided by the average of the 10-min baseline (set to 100%) prior to TBS to determine the amount of potentiation. For time course graphs, normalized fEPSP slope values were averaged and plotted as the percent change from baseline.

#### Immunohistochemistry

Mice were anesthetized and transcardially perfused with ice-cold PBS followed by 4% paraformaldehyde (PFA). Whole brains were removed and post-fixed in 4% PFA at 4° C overnight. After 24hrs, brains were transferred to a glycerol and Sorenson’s Buffer cryoprotection solution for long term storage. Frozen coronal sections were made using a sliding microtome, and slices placed in cryostorage solution containing phosphate buffer, ethylene glycol, polyvinylpyrolidine, and sucrose and stored at 4° C until staining was performed. Free floating sections were washed for 15min (3X) in PBS at RT then blocked and permeabilized (5% BSA, 5% NGS, 0.5% Triton X-100 and 1X PBS) at RT for 5-6hrs on a rocker. Slices were incubated with gephyrin (1:500 Synaptic Systems Mouse-147011) and VGAT (1:1000 Synaptic Systems Rabbit – 131003) antibodies in permeabilization solution overnight at 4°C on a rocker. Slices are washed for 20min in PBS (4X) and incubated with appropriate secondary antibodies (1:1000 ThermoFisher, Alexa-Fluor 488 and 568) for 2hrs in blocking solution. Prior to mounting with ProLong Gold, slices were washed for 20min in PBS (4X).

#### Immunocytochemistry

Coverslips containing neuronal cultures were fixed in a 4% PFA solution consisting of 4% sucrose, 1X PBS, and 50mM HEPES (pH 7.4) for 5min at RT. After fixation, the cells were blocked in a solution containing 5% BSA, 2% Normal Goat Serum (NGS), and 1X PBS at RT for 30min. Staining for surface GABA_A_R-γ2 subunit (1:500, Synaptic Systems, guinea pig – 224004) was performed under non-permeabilized conditions in the blocking solution for 1hr at RT. Following the primary antibody incubation, coverslips were washed three times for 5min each with 1X PBS. Subsequently, permeabilization was carried out using 0.5% NP-40 for 2min, followed by blocking at RT for 30min. Staining for gephyrin (1:600, Synaptic Systems, mouse, 3B11 clone, 147111) and VGAT (1:1000, Synaptic Systems, rabbit, 131003) was performed in the blocking solution for 1hr, following by three 5min washes with PBS. The coverslips were then incubated with appropriate secondary antibodies (1:1000, ThermoFisher, Alexa-Fluor 488, 568, and 547). Coverslips were washed three times for 5min and were then mounted on microscope slides using ProLong Gold mounting media (ThermoFisher).

#### Oxygen glucose deprivation (OGD) in neuronal culture

OGD was induced in DIV13-15 hippocampal neuronal cultures using HEPES-buffered solution. The OGD solution contained (in mM): 25 HEPES (pH 7.4), 140 NaCl, 5 KCl, 2 CaCl_2_, 1 MgCl_2_ and 10 sucrose (or supplemented with 10mM glucose for control conditions). To create an oxygen-deprived environment, the OGD-HEPES solution was placed in an anaerobic workstation at 37°C with a controlled atmosphere of 95% N_2_ and 5% CO_2_ (Bugbox Plus, Baker Co). Following 20min of OGD, reperfusion was initiated by replacing the HEPES solution with conditioned media and placing coverslips in an aerobic incubator for 95hrs. Neurons were then treated with various inhibitors for 1hr prior to fixation. The following inhibitors and concentrations were used (in μM): 20 CTZ, 2 tatM2NX, 2 tatScr, 5 tatCN19o, 5 KN93. In the control, no OGD condition, coverslips were fixed 96hrs following OGD-reperfusion.

#### Protein fractionation and Western immunoblotting

Mice were anesthetized, followed by rapid decapitation and brain removal. The whole hippocampus was dissected and flash frozen and stored at -80°C until membrane fractionation was performed. Membrane fraction preparation was performed as described previously(Deng et al., 2017). Whole hippocampus was homogenized in ice cold homogenization buffer containing (in mM) 10 Tris (pH7.4), 320 sucrose, 1µM EDTA, 1µM EGTA, phosphatase and protease inhibitors (ThermoScientific) using glass homogenizers, and a drill fitted with pestle. Homogenates were then transferred to 1.5 mL Eppendorf tubes and centrifuged for 10 min at 1,000 x *g* at 4°C. The supernatant was collected and placed in a clean Eppendorf tube and centrifuged at 10,000 x *g* at 4°C. The supernatant was then removed and the resulting pellet (P2) containing the membrane fraction was re-suspended in 60 µl Neuronal protein extraction reagent (ThermoScientific). Protein concentrations were quantified using Pierce BCA Protein Assay Kit (ThermoScientific) before diluting each sample in 5x SDS loading buffer to equal concentrations, and heating to 95°C for 5 minutes. Protein separation was achieved by SDS-PAGE. Western blot was performed by transferring proteins to PVDF membranes and using the following primary antibodies: CaMKII (1:2000; Cbα2 antibody, gift from KU Bayer), phosphoCaMKII T286 (1:1000; Phosphosolutions cat# p1005-286), HRP conjugated βactin (1:1000; Sigma-Aldrich), and species appropriate secondaries (anti mouse 1:5000, Jackson ImmunoResearch; anti rabbit, 1:5000; ThermoScientific).

#### Quantitative Real-time PCR

For measurement of GABA_A_R subunit transcripts, CA1 isolates were harvested 7 days following sham and CA/CPR surgeries. RNA isolations and PCR was performed as previously described(Dietz et al., 2019). Briefly, per the manufacturer’s instructions, RNA was isolated using the RNAqueous-4 PCR kit (Ambion). Approximately 0.5mg of tissue was lysed in lysis buffer and total RNA was isolated and eluted from a column with 50μL RNase-free elution buffer and further treated with Turbo DNase (Ambion). RNA (500ng) was reverse transcribed to single stranded cDNA using the iScript cDNA Synthesis Kit (BioRad). Real-time PCR reactions using ssoFast PCR mastermix (Biorad) were performed on BioRad CFX connect detection system and performed in triplicate using 50ng of cDNA. Taqman (ThermoScientific) primers used to detect *GABRG2*, *GABRB3*, *GABRA1* and 18s transcripts. Cycle parameters used were 95°C for 10min followed by 40 cycles of 95°C for 15s and 60°C for 30s. Relative expression levels were calculated using ΔΔCT as the ratio of the target gene to the housekeeping gene18s.

### Image Acquisition and Data Analysis

#### Confocal microscopy

Confocal images were acquired on a Zeiss Axio Observer.Z1 upright microscope equipped with a Yokogawa CSU-X1 spinning disk unit; a 63X oil immersion objective using 2x digital zoom (Plan-Apo/1.4 NA); an Evolve 512 EM-CCD camera (Photometrics) with 16-bit range; and SlideBook 6.0. Alternatively, neurons were imaged using an Olympus FV1000 laser scanning confocal microscope, 60x oil immersion objective with 2x digital zoom and Fluoview software (Olympus FluoView, FV10-ASW). Images on both microscopes were attained at 0.3 μm intervals (4μm Z-stack projection). Cluster analysis was performed using ImageJ (NIH) by selecting regions of interest (ROIs) to differentiate between dendritic and somatic compartments. A user-based threshold was determined by sampling several images per condition across all conditions and clusters were defined as a minimum size of 0.05μm^2^. For IHC experiments, CA1 hippocampus was identified by VGAT staining of the pyramidal cell layer (PCL). ROIs captured both the PCL and *Stratum radiatum* in the same frame, and different user-based thresholds were used for the cell bodies and dendrites. Density was calculated by dividing the number of clusters by the ROI area (per μm^2^). A minimum of 5 animals per condition were utilized, with two slices per animal analyzed. Analysis was performed blind to surgical condition. For the *in vitro* experiments, ROIs were delineated by tracing along dendrites. Density of clusters was calculated by measuring the length of the delineated dendrites (per 10μm). A total of 30-36 neurons were analyzed per condition from three independent hippocampal preparations.

#### Statistical Analysis

Data are presented as mean ± SEM. Statistical significance was determined using appropriate statistical tests as indicated in the figure legends. A p-value ≤ 0.05 was used to declare significance. All statistical analyses were performed on GraphPad Prism v9.4.

## RESULTS

### E/I balance is disrupted following CA/CPR

To assess the effect of GCI on E/I balance, we employed patch-clamp electrophysiology to record evoked GABA (0mV) and AMPA (-70mV) responses from CA1 pyramidal neurons 7 days following CA/CPR or sham surgery (Fig 1A), a post-acute timepoint at which cell death cell death processes have subsided and perturbations in synaptic function are reflective of the surviving network. The ratio of IPSC (GABA) to EPSC (AMPA) amplitude was greater in CA/CPR compared to sham, indicating increased synaptic inhibitory function relative to excitatory function. This increase in inhibitory function following CA/CPR was observed despite a decrease in release probability in the CA/CPR group compared to sham as measured by the paired pulse ratio of IPSCs, (Fig S1A), with no differences in EPSC release probability observed (Fig S1B). This suggests a potential enhancement of postsynaptic inhibitory function which may outweigh a reduced presynaptic function, leading to an overall reduction in the E/I ratio. To determine the effect of increased inhibition on LTP deficits following CA/CPR, we performed extracellular field recordings in the CA1-Schaffer collateral pathway. We then treated hippocampal sections from CA/CPR mice with picrotoxin (5μM) (Costa and Grybko, 2005), a GABA_A_R pore blocker, to test the effect of GABAergic inhibition on LTP. Following theta burst stimulation (TBS), CA/CPR slices without treatment exhibited impaired LTP, consistent with previous reports (Dietz et al., 2019; Orfila et al., 2018; Orfila et al., 2014). However, treatment with picrotoxin restored LTP to sham levels (Fig 1C-E). No differences in the input-output curve (Fig S1E) and paired pulse ratio (Fig S1F) were observed in the field recordings, suggesting enhanced postsynaptic GABA_A_R function contributes to LTP deficits at post-acute timepoints following CA/CPR.

**Figure 1.**
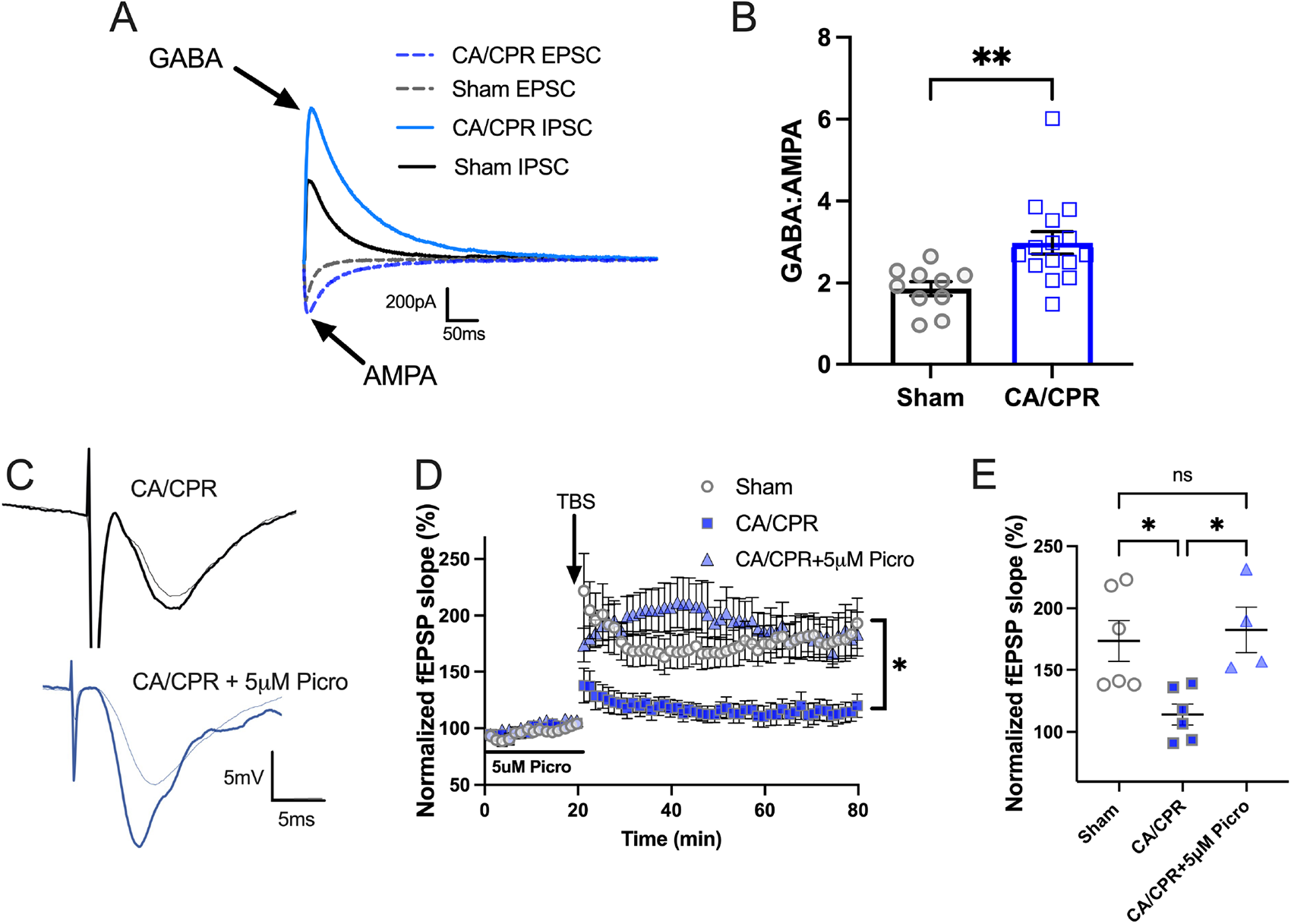
Excitatory/Inhibitory balance is disrupted following CA/CPR. A. Shown are representative traces for evoked EPSCs (solid) and IPSCs (dotted) recorded from the same CA1 hippocampal neurons of CA/CPR (blue) and sham (gray/black) mice. The AMPA eEPSCs are recorded with the neuron held at -70 mV and GABA eIPSCs recorded at 0 mV. B. Quantification of the absolute value of the ratio of GABA eIPSC amplitude to the AMPA eEPSC amplitudes; n=11-15 cells/ 3-5 animals per condition; unpaired t-test. C. Representative fEPSP traces from the *stratum radiatum* region of the CA1-hippocampus in CA/CPR slices without treatment (black) and CA/CPR slices (blue) treated with 5µM picrotoxin during baseline. D. LTP data presented as a percentage of the baseline, where baseline is set at 100%. E. Normalized slope of fEPSPs after theta burst stimulus; n=4-6 slices/ 4-5 animals per condition; one-way ANOVA, Tukey’s post hoc test. Values represent mean ± SEM. *p<0.05; ** p<0.01.

### GABAergic sIPSC amplitude is increased following CA/CPR and can be rapidly reduced following TRPM2 ion channel inhibition

To determine whether the increase in inhibitory function is pre or postsynaptic, we recorded spontaneous IPSCs (sIPSCs) from CA1 neurons in CA/CPR and sham operated mice and assessed amplitude, frequency, and decay kinetics. The cumulative frequency of sIPSCs amplitude was shifted rightward in CA/CPR mice compared to sham operated mice, indicating an overall increase in amplitude (Fig 2B). Consistent with this, CA/CPR mice exhibited increased mean sIPSC amplitude compared to sham (Fig 2A-C_1_). No differences in mean frequency nor tau decay were detected (Fig 2C_2_-2C_3_), suggesting the increase in inhibitory function is primarily postsynaptic.

Given the potential link between elevated GABAergic inhibition and LTP impairments in our experiments, we next asked whether TRPM2 inhibition, previously shown to restore LTP after CA/CPR (Dietz et al., 2019; Dietz et al., 2021), potentially acts through postsynaptic GABAergic changes. To test this, we measured sIPSCs from CA1 neurons and bath applied tatM2NX (2μM), a potent and specific TRPM2 inhibitor(Cruz-Torres et al., 2020), for at least 20 minutes prior to recording. CA/CPR slices treated with tatM2NX showed a reduction in sIPSC amplitude compared to CA/CPR slices without treatment, shifting the cumulative frequency of sIPSCs amplitude leftward, (Fig 2B). The mean amplitude was also reduced in tatM2NX treated CA/CPR slices compared to slices without treatment (Fig 2C_1_). No differences in mean frequency nor tau decay were detected across all conditions (Fig 2C_2_-2C_3_).

We then wanted to determine whether TRPM2 inhibition can *rapidly* reverse the CA/CPR-induced increase in amplitude within the same cell. We patched CA1 neurons from sham and CA/CPR mice and measured sIPSC amplitude, frequency and tau decay before and after bath application of clotrimazole (CTZ, 20μM), a fast-acting TRPM2 pore blocker(Hill et al., 2004). CA/CPR slices treated with CTZ shifted the cumulative frequency of sIPSC amplitude leftward (Fig 2E), indicating CTZ treatment restored sIPSC amplitude to sham levels. TRPM2 inhibition by CTZ rapidly reduced the CA/CPR-induced increase in sIPSC mean amplitude (Fig 2F_1_). Surprisingly, we observed a reduction in mean frequency in CA/CPR slices following bath application of CTZ (Fig 2F_2_). This was likely an off-target effect as the more specific TRPM2 inhibitor, tatM2NX, had no effect on frequency. However, consistent with the tatM2NX data, no differences were detected in tau decay across all conditions (Fig 2F_3_). Taken together, these data suggest the increase in GABAergic function observed following CA/CPR is primarily postsynaptic and can be rapidly reversed following TRPM2 ion channel inhibition.

**Figure 2.**
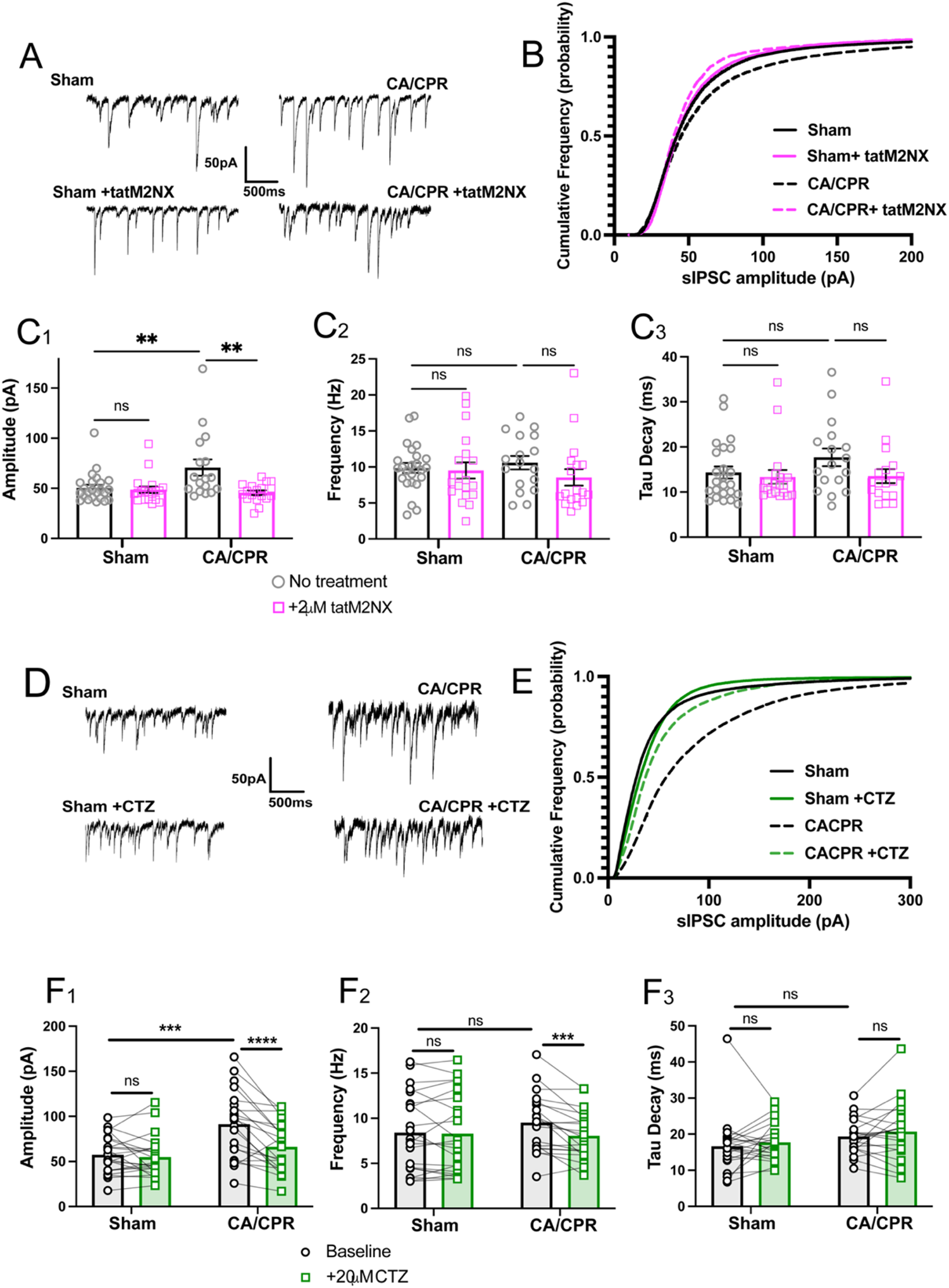
GABAergic sIPSC amplitude is increased following CA/CPR and can be rapidly reduced following TRPM2 ion channel inhibition. A. Representative traces from sham (top, left), sham with 2µM tatM2NX bath applied (bottom, left), CA/CPR (top, right), CA/CPR with bath applied tatM2NX (bottom right) of whole-cell voltage clamp recordings of sIPSC events recorded from CA1 pyramidal neurons in acute hippocampal slices using CsCl based internal solution holding at -60mV. B. Cumulative frequency distribution of sIPSC amplitude from sham cells (black, solid), sham+ tatM2NX (black, dotted), CA/CPR (pink, solid), CA/CPR+ tatM2NX (pink, dotted). C. Mean amplitude (1), frequency (2), and tau decay (3) kinetics were measured from sIPSC events in different cells from sham and CA/CPR operated mice with and without bath application of tatM2NX; n=18-24 cells/ 8-10 animals per condition; One-Way ANOVA, Tukey’s posthoc test. D. Representative traces from sham (top, left), sham after clotrimazole (CTZ) from the same cell (bottom, left), CA/CPR (top, right), CA/CPR after CTZ from the same cell (bottom, right). E. Cumulative frequency distribution of sIPSC amplitude from CA/CPR (black, dotted) following treatment with CTZ (20μM) in the same cell (green, dotted). F. Mean amplitude (1), frequency (2), and tau decay (3) kinetics were measured from sIPSC events in the same cell before and after bath application of CTZ from sham and CA/CPR operated mice; n=21-22 cells/ 9-11 animals per condition; Two-Way ANOVA with Repeated Measures, Sidak’s post hoc test. Values represent mean ± SEM. *p<0.05; ** p<0.01, ***p<0.001, ****p<0.0001.

### CA/CPR induces a persistent increase in clustering and density of gephyrin

We next investigated the mechanisms underlying the increase in inhibitory synaptic function driven by TRPM2 following CA/CPR. The increase in sIPSC amplitude suggests more postsynaptic receptor clustering at the synapse and a larger postsynaptic domain. To test this, we performed immunohistochemistry 7 days following sham and CA/CPR procedures and stained for the postsynaptic inhibitory scaffold protein, gephyrin, and the presynaptic inhibitory synapse marker, vesicular GABA transporter (VGAT). There was no difference in VGAT cluster area nor density between CA/CPR and sham in either the *stratum radiatum* or the *stratum pyramidale* (Fig 3C-D). In contrast, we observed an increase in gephyrin cluster density in the *stratum radiatum* (Fig 3C). In the *stratum pyramidale*, both gephyrin cluster area and density were increased in CA/CPR mice compared to sham (Fig 3D). These changes in postsynaptic GABAergic components were not due to altered gephyrin protein expression (Fig 2SA-B), nor altered transcription of various GABA_A_R subunits (Fig 2SC-E). These findings align well with our electrophysiology data, suggesting CA/CPR enhances postsynaptic GABAergic function through increased receptor density, which can be rapidly reversed without affecting presynaptic inhibitory function.

**Figure 3.**
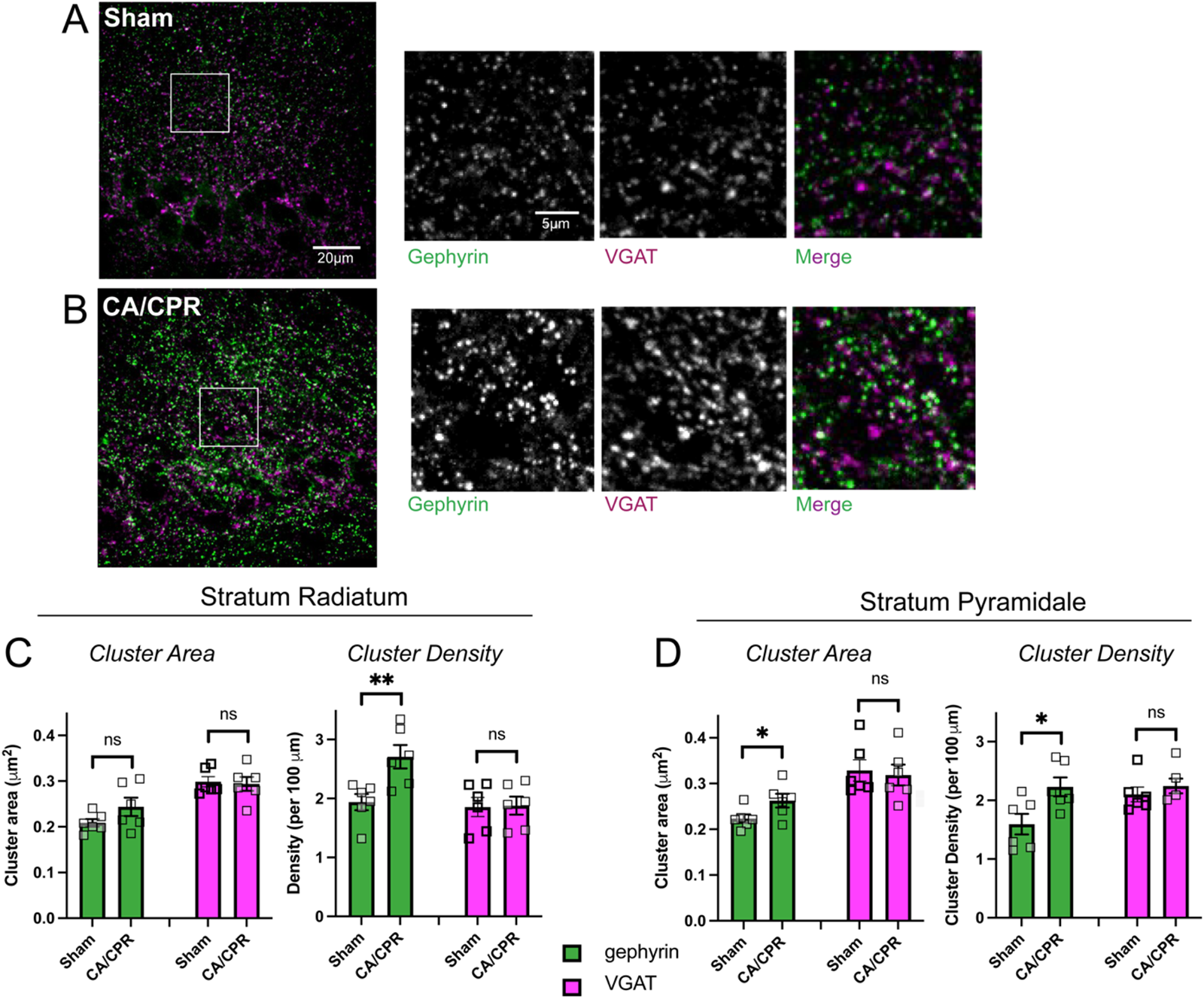
CA/CPR induces a persistent increase in gephyrin clustering and density. A-B. Shown are representative images of the CA1 region of the hippocampus from (A) sham operated and (B) CA/CPR mice. Gephyrin staining is shown in green and VGAT is shown in magenta. C. Quantification of the cluster area and density of both gephyrin and VGAT in the *stratum radiatum*; n=6 animals per group; unpaired t-test. D. Quantification of the cluster area and density of both gephyrin and VGAT in the *stratum pyramidale*; n= 6 animals per group; unpaired t-test. Values represent mean ± SEM. *p<0.05; ** p<0.01.

### OGD induces a persistent increase in the clustering and density of postsynaptic GABAergic proteins

To directly visualize GABA_A_Rs and interrogate the mechanisms contributing to the enhancement of synaptic GABA_A_R clustering, we extended our studies to a well-established *in vitro* model of GCI (Arancibia-Carcamo et al., 2009; Garcia et al., 2021; Smith et al., 2017). We exposed dissociated hippocampal neurons to 20-minute oxygen-glucose deprivation (OGD) followed by reperfusion and fixed the neurons at varying timepoints to mimic the delayed 7-day timepoint we analyzed post-CA/CPR (Fig 4A). To assess inhibitory synaptic size and postsynaptic receptor clustering, we immunostained for gephyrin, surface GABA_A_Rs (γ2 subunit), and VGAT, and evaluated cluster area and density of these markers in dendrites of pyramidal neurons using confocal microscopy (Fig 4B). We observed a decrease in cluster area and density of all synaptic GABAergic components 24hrs post-OGD, consistent with prior work showing an acute loss of GABAergic synapses (Garcia et al., 2021). However, by 48hrs, these measurements were no different from control levels. By 72- and 96hrs, both the cluster area (Fig 4C-E) and density (Fig 4F-H) increased for all synaptic GABAergic components. The increase in the presynaptic marker, VGAT, was unexpected given the *in vivo* CA/CPR data showed no changes in VGAT clustering (Fig 3C-D) nor a presynaptic functional effect (Fig 2). Despite this discrepancy, both postsynaptic markers exhibited an acute reduction in clustering followed by an increase in the chronic phase, consistent with the *in vivo* results shown prior (Fig 3). Thus, we utilized this *in vitro* system, specifically the 96hr timepoint, to investigate the mechanism of TRPM2-mediated postsynaptic GABAergic enhancement.

**Figure 4.**
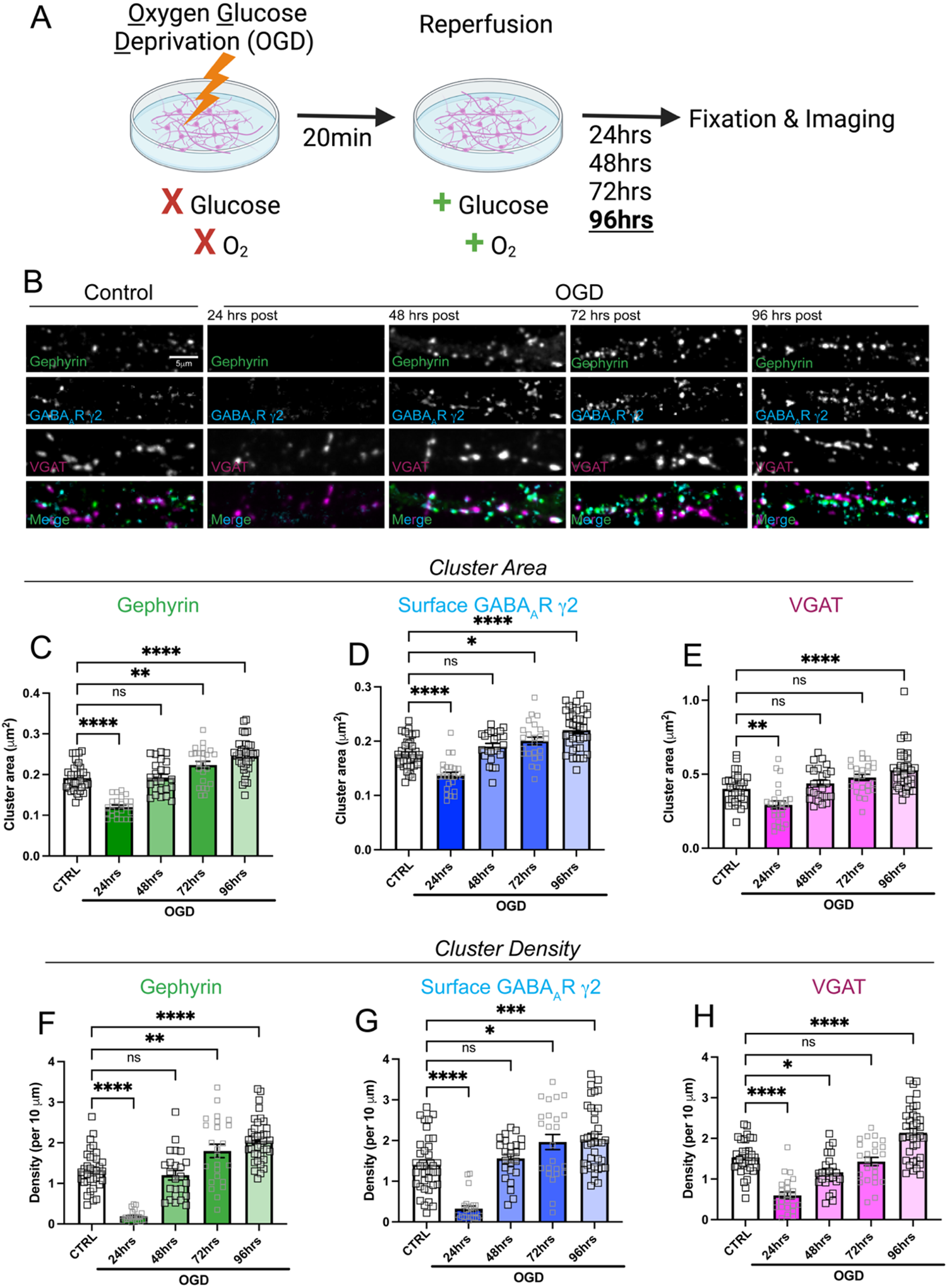
OGD induces a persistent increase in clustering and density of postsynaptic GABAergic proteins. A. Cartoon illustrating timeline for *in vitro* OGD experiments. Dissociated hippocampal neurons were reperfused following 20min OGD exposure. Neurons were fixed 24-, 48-, 72-, and 96hrs following reperfusion. B. Representative confocal images of dendritic segments from pyramidal neurons stained for gephyrin (green), GABA_A_R-γ2 subunit (cyan), VGAT (magenta). C-E. Quantification of cluster area for (C) gephyrin, (D) surface GABA_A_R-γ2, and (E) VGAT; n=30-36 neurons per condition; One-Way ANOVA, Dunnett’s post hoc. F-H. Quantification of cluster density for (F) gephyrin, (G) surface GABA_A_R-γ2, and (H) VGAT; n=30-36 neurons per condition; One-Way ANOVA, Dunnett’s post hoc. Values represent mean ± SEM. *p<0.05; ** p<0.01, ***p<0.001, ****p<0.0001.

### The TRPM2 ion channel mediates the OGD-induced increase in clustering of postsynaptic GABAergic components

We next investigated whether TRPM2 activation affects the density of postsynaptic GABAergic proteins following OGD. We subjected hippocampal neurons to 20min OGD-reperfusion and treated with either tatM2NX or CTZ 1hr prior to fixation (Fig 5A). Treatment with tatM2NX, a potent and specific TRPM2 inhibitor, reduced the OGD-induced increase in gephyrin (Fig 5C) and surface GABA_A_R-γ2 subunit cluster area (Fig 5D) as well as the presynaptic marker, VGAT (Fig 5E). We then performed similar experiments using the noncompetitive TRPM2 blocker, CTZ, and found CTZ treatment also significantly reduces the OGD effect on gephyrin and surface GABA_A_R-γ2 cluster area with no effect on VGAT across all conditions (Fig 5H). Consistent with these results, both tatM2NX and CTZ treatment reduced the cluster density of all synaptic GABAergic proteins (Fig S3). Using two pharmacological inhibitors of TRPM2, our data strongly support that the persistent OGD-induced increase in postsynaptic GABA_A_R density is TRPM2-dependent, consistent with the *in vivo* functional data which implicates TRPM2 activity in the increase in GABAergic sIPSC amplitude post-CA/CPR (Fig 2).

**Figure 5.**
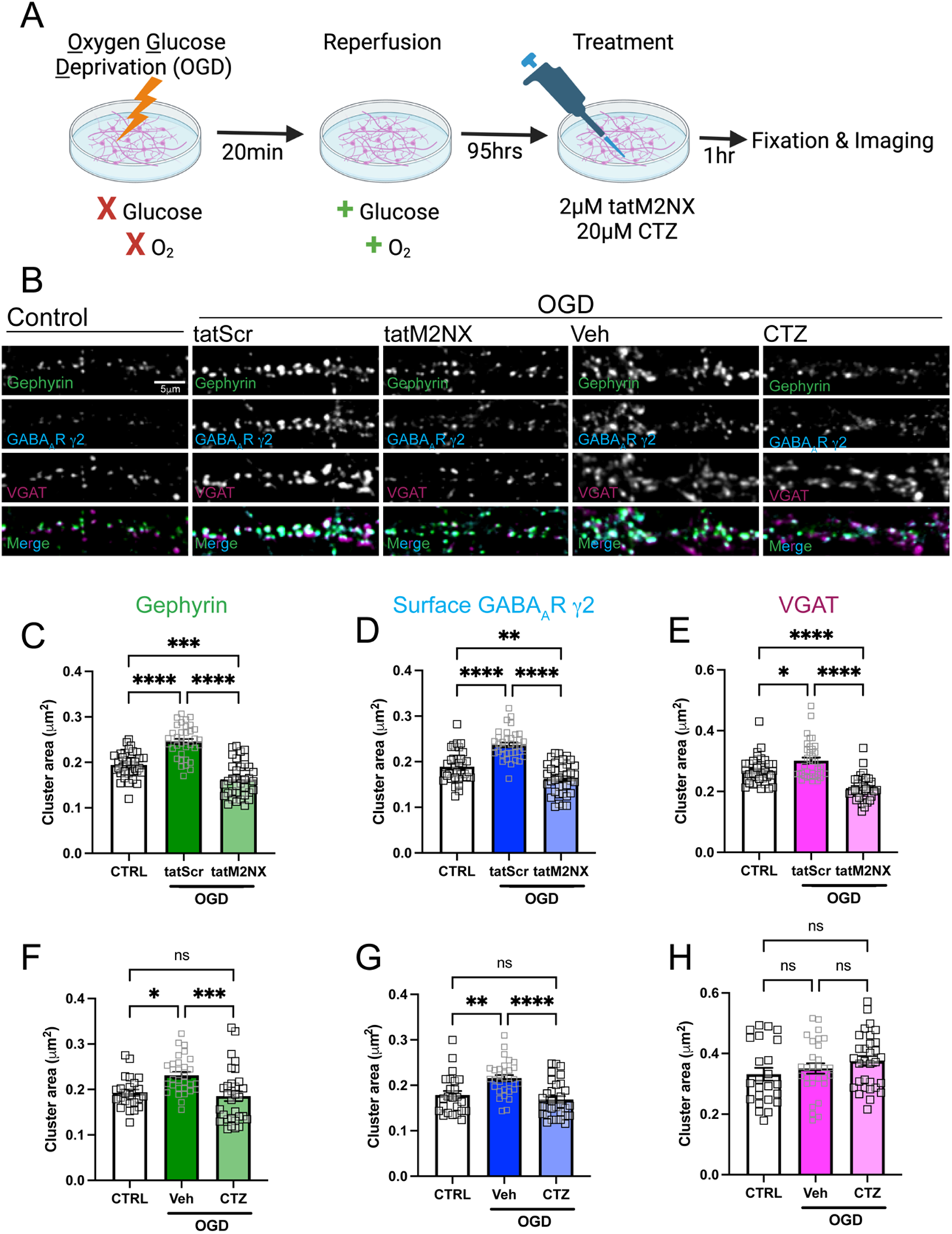
The TRPM2 ion channel mediates the OGD-induced increase in clustering of postsynaptic GABAergic proteins. A. Cartoon illustrating experimental timeline for neurons subjected to OGD-reperfusion and treated with tatM2NX (2μM) or CTZ (20μM) 1hr prior to fixation. B. Representative confocal images of dendritic segments from pyramidal neurons immunostained for gephyrin (green), GABA_A_R-γ2 subunit (cyan), VGAT (magenta). C-E. Quantification of cluster area following treatment with tatM2NX for (C) gephyrin, (D) surface GABA_A_R-γ2, and (E) VGAT; n=30-36 neurons per condition; One-Way ANOVA, Tukey’s posthoc. F-H. Quantification of cluster area following treatment with CTZ for (F) gephyrin, (G) surface GABA_A_R-γ2, and (H) VGAT, n=30-36 neurons per condition; One-Way ANOVA, Tukey’s posthoc. Values represent mean ± SEM. *p<0.05; ** p<0.01, ***p<0.001, ****p<0.0001.

### Ca^2+^ signaling and CaMKII activity contributes to CA/CPR-induced increase in GABAergic sIPSC amplitude

While these data implicate TRPM2 in the ischemia-induced enhancement of postsynaptic inhibitory function, the downstream signaling required to regulate GABA_A_R density remains yet to be determined. We hypothesized TRPM2-mediated Ca^2+^ influx may modulate postsynaptic GABA_A_R function. To examine this, we performed patch-clamp recordings with a high concentration of BAPTA (10mM), a selective Ca^2+^ buffer, in the patch pipette and recorded from the same cell before and after CTZ treatment. Our results showed Ca^2+^ chelation in the postsynaptic neuron occluded the CA/CPR-induced enhancement of sIPSC mean amplitude, as there was no longer a difference between CA/CPR and sham conditions (Fig 6B_1_). Further, we did not observe a change in amplitude following CTZ treatment in cells recorded with BAPTA in the patch pipette (Fig 6B_1_). We also found no differences between the CA/CPR and sham conditions and the CTZ treatment in the frequency nor the tau decay of sIPSCs when BAPTA was present (Fig 6B_2-3_). Given there was no additive effect of BAPTA and CTZ on sIPSC amplitude, this suggests that postsynaptic Ca^2+^ signaling mediates the observed TRPM2-dependent increase in inhibition.

CaMKII is a Ca^2+^-dependent kinase that has been previously shown to be required for activity-dependent postsynaptic GABA_A_R potentiation and clustering (Chiu et al., 2018; Cook et al., 2021; Cook et al., 2022; Petrini et al., 2014). We therefore hypothesized TRPM2-mediated Ca^2+^ signaling might activate CaMKII to enhance postsynaptic GABAergic function. To test this, we used western immunoblotting to detect total protein levels of CaMKII in the membrane fractions of whole hippocampus from CA/CPR and sham mice 7 days following the surgeries. Our data show that CaMKII levels are significantly reduced in the membrane fraction of whole hippocampi (Fig 6C). Interestingly, we observed a significantly elevated level of T286 phosphorylation in the synaptic membrane fraction obtained from CA/CPR mice compared to sham (Fig 6D), indicating a sustained increased synaptic CaMKII activity following GCI.

To further examine the role of elevated CaMKII activity on postsynaptic GABAergic function following CA/CPR, we used patch-clamp electrophysiology to record sIPSCs while inhibiting postsynaptic CaMKII using tatCN19o (5μM) in the patch pipette. Our results show that inhibition of postsynaptic CaMKII activity occluded the CA/CPR-induced increase in sIPSC amplitude, as indicated by equivalent sIPSC amplitudes observed in tatCN19o treated cells and sham conditions (Fig 5F_1_). Additionally, CTZ no longer impacted the mean amplitude of sIPSCs in the CA/CPR mice recorded with tatCN19o (Fig 5F_1_). Similar to the effects observed with Ca^2+^ chelation, we did not observe a change between the CA/CPR and sham conditions and the CTZ treatment in the frequency and tau decay of sIPSCs when the CaMKII inhibitor was present (Fig 2F_2-3_). These results indicate that CaMKII-dependent increases in postsynaptic inhibitory function are likely downstream of Ca^2+^ entry via TRPM2 channel activity.

**Figure 6.**
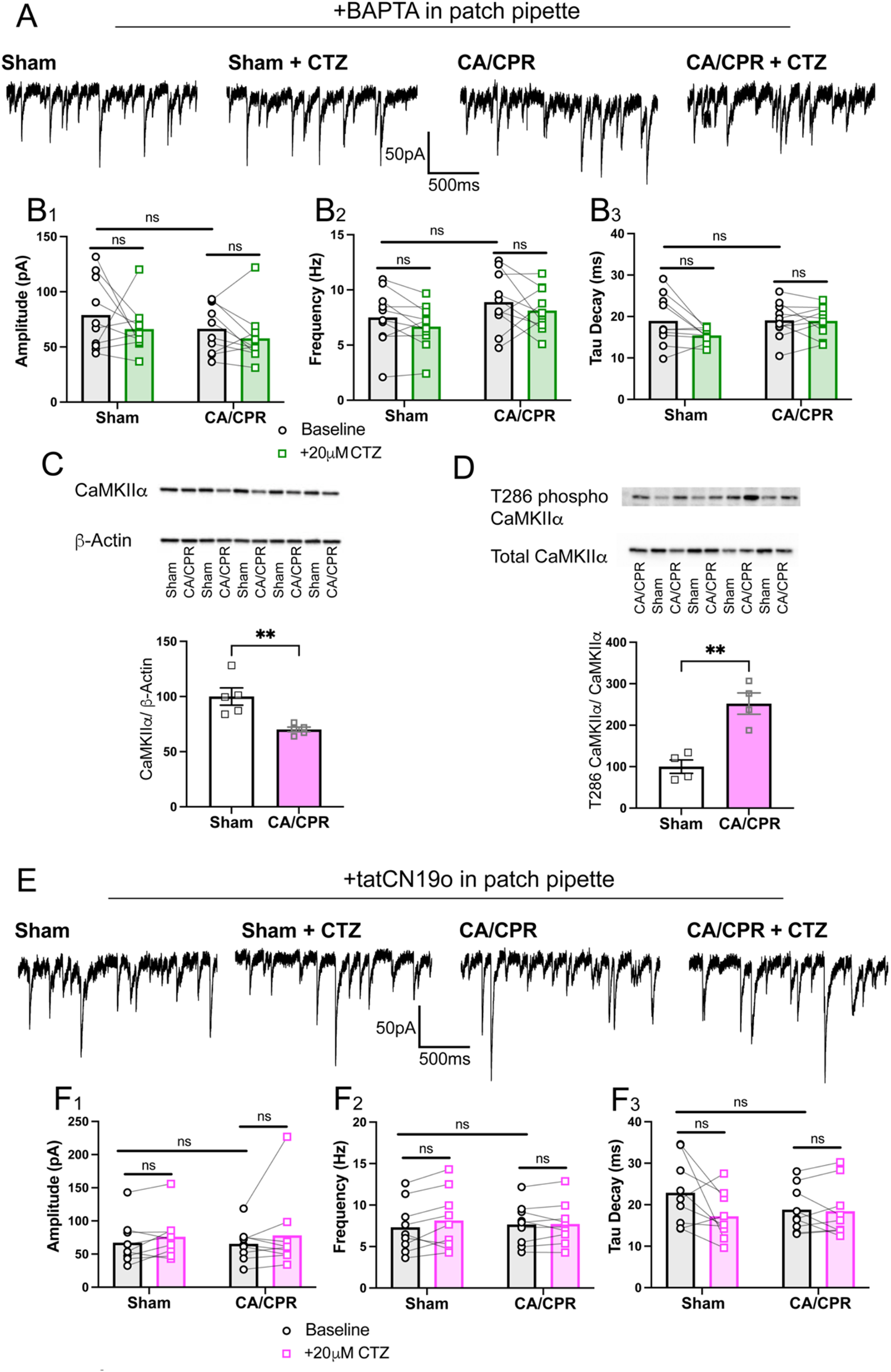
Ca^2+^ signaling and CaMKII activity contributes to CA/CPR-induced increase in amplitude of GABAergic sIPSCs. A. Representative traces from sham (far left), sham after CTZ from the same cell (center left), CA/CPR (center right), CA/CPR after CTZ from the same cell (far right). BAPTA (10mM) was included in the patch pipette across all conditions. B. Mean amplitude (1), frequency (2), and tau decay (3) kinetics were measured from sIPSC events in the same cell before and after bath application of CTZ from sham and CA/CPR operated mice with 10mM BAPTA included in patch pipette; n= 10 cells/ 4-7 animals per condition; Two-Way ANOVA with Repeated Measures, Sidak’s post hoc test. C. Western immunoblot measuring total levels of CaMKII normalized to βactin loading control in membrane fraction of whole hippocampi from sham and CA/CPR operated mice; n=5 animals per condition; unpaired t-test. D. Western immunoblot measuring levels of T286 phosphorylation of CamKII normalized to total CaMKII in the membrane fraction of whole hippocampi from sham and CA/CPR operated mice. Grubbs’ outlier test was used for exclusion of one higher, outlier value in the CA/CPR group; n=4 animals per condition; unpaired t-test. E. Representative traces from sham (far left), sham after CTZ from the same cell (center left), CA/CPR (center right), CA/CPR after CTZ from the same cell (far right). TatCN19o (5μM) was included in the patch pipette across all conditions. F. Mean amplitude (1), frequency (2), and tau decay (3) kinetics were measured from sIPSC events in the same cell before and after bath application of CTZ from sham and CA/CPR operated mice with 5μM tatCN19o included in patch pipette; n=9 cells/ 4-5 animals per condition; Two-Way ANOVA with Repeated Measures, Sidak’s post hoc test. Values represent mean ± SEM. *p<0.05; ** p<0.01.

### Ca^2+^-dependent CaMKII activity contributes to increased clustering of postsynaptic GABAergic components via a TRPM2-dependent pathway

We next investigated whether TRPM2-and CaMKII-mediated enhancement of postsynaptic sIPSC amplitude was due to an increase in the density of postsynaptic GABA_A_Rs. To assess this, we used the 96hr timepoint following OGD-reperfusion in dissociated rat hippocampal neurons and treated cells with vehicle (control), tatCN19o alone, or combined tatCN19o and tatM2NX 1hr prior to fixation and immunostained for GABAergic synaptic markers (Fig 7A). We found that treatment with tatCN19o reduced the cluster area of the GABA_A_R-γ2 and gephyrin following OGD compared to the OGD vehicle control (Fig 7B-C). Additionally, we observed a significant decrease in the cluster area of the postsynaptic components in the combined tatM2NX and tatCN19o condition compared to the control (veh) condition following OGD; however, there was no additional reduction in the cluster size of the postsynaptic inhibitory synaptic proteins in the combined treatment condition compared to the tatCN19o alone (Fig 7B-C). In agreement with this, tatCN19o alone and combined treatment tatM2NX occluded the OGD-induced increase in cluster density of postsynaptic GABAergic components (Fig S4A-B). We found no differences across all conditions when measuring VGAT cluster area (Fig 7E). These results suggest that TRPM2 and CaMKII likely converge on the same pathway to regulate postsynaptic GABA_A_R density.

The tatCN19o peptide was shown to block both the Ca^2+^-independent autonomous and Ca^2+^-stimulated CaMKII activity (Vest et al., 2010). Based on our functional data showing Ca^2+^ signaling was required for the increase in sIPSC amplitude and is reversible with Ca^2+^ chelation (Fig 6), we hypothesized that the Ca^2+^-stimulated CaMKII activity contributes to elevated GABA_A_R clustering. To test this, we treated neurons 95hrs following OGD-reperfusion with KN93 (5μM), a small molecule CaMKII inhibitor known to preferentially block Ca^2+^-stimulated CaMKII activity (Fig 7A) (Vest et al., 2010). We found that KN93 significantly reduced the cluster area of both postsynaptic GABA proteins following OGD compared to the OGD vehicle control condition (Fig 7F-G). This reduction in cluster area of postsynaptic components persisted in the combined KN93 and tatM2NX condition, showing no differences compared to KN93 treatment alone following OGD (Fig 7F-G). Similarly, both KN93 and combined conditions reduced the OGD-induced increase in cluster density of gephyrin and GABA_A_R-γ2 (Fig S4D-E). While there was a significant increase in the presynaptic marker, VGAT, cluster area in the vehicle OGD condition compared to the control no OGD group, neither KN93 nor the combined KN93 and tatM2NX treatment significantly affected VGAT cluster size (Fig 7H). Altogether, these data suggest that the TRPM2 channel and Ca^2+^-stimulated CaMKII activity converge on the same molecular pathway to regulate postsynaptic GABA_A_ receptor density at delayed timepoints following OGD.

**Figure 7.**
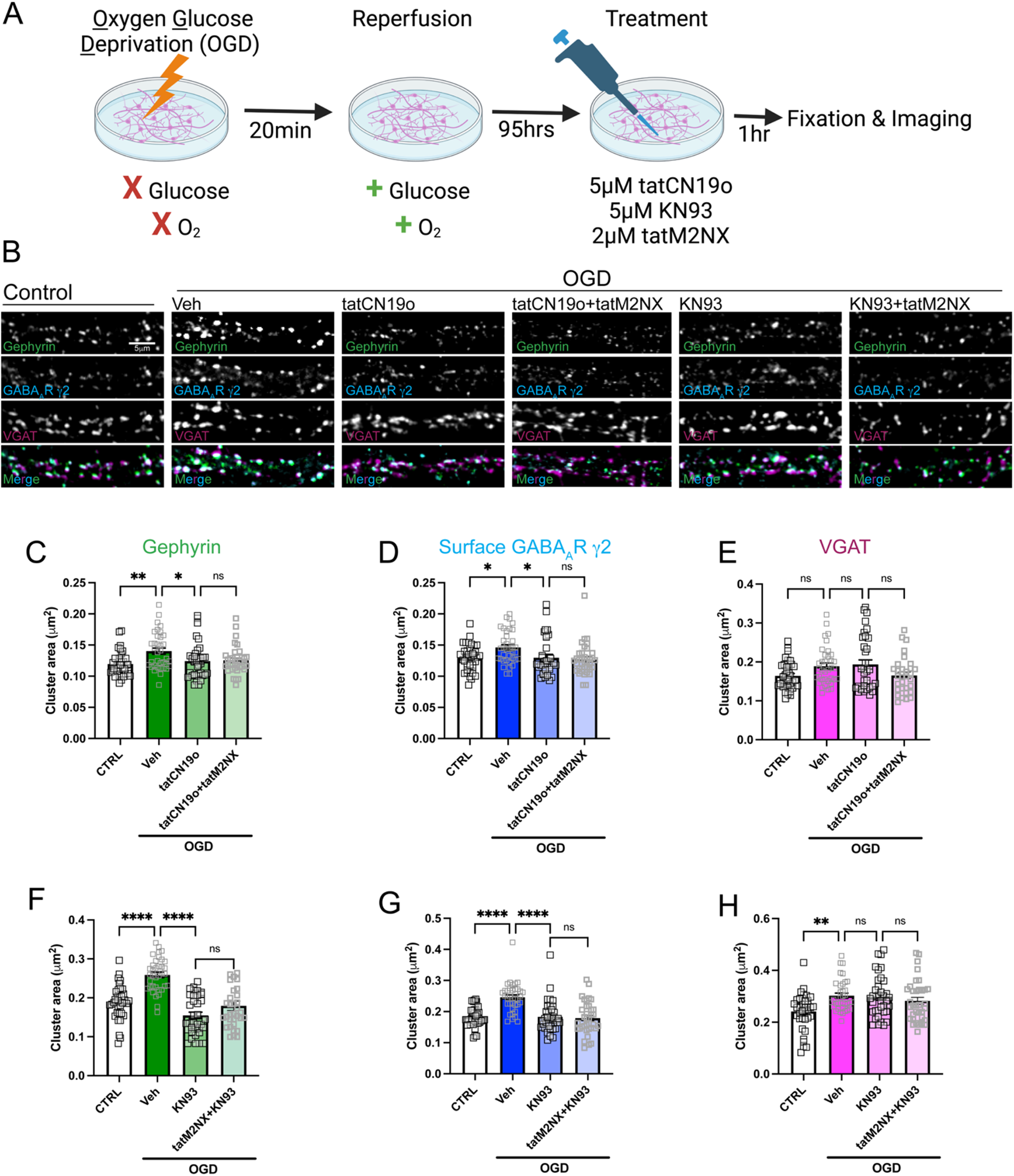
Ca^2+^-dependent CaMKII activity contributes to the increased clustering of postsynaptic GABAergic proteins via a TRPM2-dependent pathway. A. Cartoon illustrating experimental timeline for neurons subjected to OGD-reperfusion and treated with tatCN19o (5μM), KN93 (5μM), or combined treatment with tatM2NX (2μM) 1hr prior to fixation. B. Representative confocal images of dendritic segments from pyramidal neurons stained for gephyrin (green), GABA_A_R-γ2 subunit (cyan), VGAT (magenta). C-E. Quantification of cluster area following treatment with tatCN19o or tatCN19o+ tatM2NX for (C) gephyrin, (D) surface GABA_A_R-γ2, and (E) VGAT; n=30-36 neurons per condition; One-Way ANOVA, Tukey’s posthoc. F-H. Quantification of cluster area following treatment with KN93 or KN93+tatM2NX for (E) gephyrin, (F) surface GABA_A_R-γ2, and (G) VGAT; n=30-36 neurons per condition; One-Way ANOVA, Tukey’s posthoc. Values represent mean ± SEM. *p<0.05; ** p<0.01, ***p<0.001, ****p<0.0001.

## DISCUSSION

Here, we combine *in vitro* and *in vivo* approaches to elucidate the prolonged enhancement of postsynaptic GABAergic function in the hippocampus, accounting for the reduction in E/I ratio and driving the LTP deficits following CA/CPR. Using an *in vitro* model that emulates the direct ischemic insult to the hippocampus observed in GCI, we found a shift from an acute reduction in GABAergic signaling to a sustained elevation in the clustering of GABAergic proteins at synaptic sites following OGD-reperfusion. To our knowledge, this is the first study to employ the OGD *in vitro* system to examine synaptic alterations days after ischemic insult following completion of cell death processes. Using this novel *in vitro* paradigm and a murine model of GCI, we identified the TRPM2 ion channel as a potential mediator of GABAergic synaptic enhancement. TRPM2 inhibition effectively blocked the CA/CPR-induced increase in sIPSC amplitude and the OGD-induced effect on cluster area of postsynaptic GABAergic components. Furthermore, our data revealed that TRPM2 and Ca^2+^-CaMKII are both required for the elevated postsynaptic GABAergic function. Chelation of Ca^2+^ and CaMKII blockade both alleviated the CA/CPR effect on sIPSC amplitude and reversed the OGD-induced increase in cluster area of synaptic GABAergic proteins. Additionally, blockade of the TRPM2 channel showed no additive effect on the reduction in sIPSC amplitude, nor GABA_A_R clustering following CaMKII inhibition, suggesting that inhibitory synaptic function is enhanced via a TRPM2-CaMKII-dependent pathway.

Elevation of excitatory signaling with concurrent loss of inhibitory in the acute period is well-defined. Excitotoxicity, characterized by aberrant NMDA and AMPA receptor activation and intracellular Ca^2+^ overload represents one of the many signaling events that contribute to neuronal cell demise (Kostandy, 2012; Lau and Tymianski, 2010; Moskowitz et al., 2010). This hyperexcitability is further exacerbated by a profound loss of inhibitory synaptic strength and density, tipping the E/I ratio in favor of excitatory signaling (Alicke and Schwartz-Bloom, 1995; Mele et al., 2014; Schwartz-Bloom and Sah, 2001). The role of GABAergic signaling beyond acute cell death and excitotoxicity has received relatively less attention; however, emerging work provides evidence of a subsequent reduction in E/I balance in the surviving network, primarily driven by enhanced inhibition through extrasynaptic (tonic) and the relatively less studied synaptic (phasic) GABA_A_R signaling (Carmichael, 2012; Joy and Carmichael, 2021). Excessive tonic inhibition has been demonstrated to hinder cortical and hippocampal recovery in animal models of focal ischemia (Clarkson et al., 2010; Lake et al., 2015; Orfila et al., 2019). Additionally, phasic inhibition, achieved through rapid activation of synaptic GABA_A_Rs, has been shown to be enhanced in the peri-infarct cortex following stroke, correlating with motor and cognitive deficits (Hiu et al., 2016). In line with these findings, our study contributes to the existing literature by describing elevated postsynaptic phasic inhibition in the hippocampus following global insult and its potential role in impairing excitatory LTP. The question of whether excessive tonic inhibition also contributes to hippocampal synaptic plasticity deficits following GCI warrants further investigation. Altogether, our observation of elevated postsynaptic (phasic) GABAergic signaling, which can be pharmacologically interrupted/ targeted without the unfavorable adverse effects of directly blocking GABA_A_R, offers a promising approach for therapeutic development.

In our study, we identify the TRPM2 ion channel as a novel mediator of inhibitory function and E/I balance following cerebral ischemia-reperfusion injury. Although TRPM2 has been extensively studied in the context of neuronal cell death, with acute inhibition of the channel conferring neuroprotection (Jia et al., 2011; Nakayama et al., 2013; Shimizu et al., 2013), recent evidence has highlighted its involvement in the post-acute phase, indicating a role for the channel in mechanisms in the expression of synaptic plasticity and its promise as a neurorestorative therapeutic target (Dietz et al., 2019; Dietz et al., 2021; Xie et al., 2011). Our prior study revealed sustained TRPM2 activity during the post-acute period following GCI has been found to contribute to ischemia-induced LTP impairment via a calcineurin-GSK3β-dependent pathway within glutamatergic synapses (Dietz et al., 2019). In contrast, our data is the first to suggest an additional role for TRPM2 in the regulation of inhibitory synaptic strength following ischemia-reperfusion and reveal Ca^2+^ and CaMKII as the most plausible downstream mediators. Consistent with this, previous evidence supports TRPM2 as the Ca^2+^ source for CaMKII activity in the context of cell cycle regulation (Cai et al., 2023; Wang et al., 2017), suggesting this may be a common mechanism across multiple cellular functions. Moreover, our data indicate that continuous Ca^2+^ stimulus is required for ongoing CaMKII activation. Notably, a prominent property of the TRPM2 channel is its ability to conduct substantial Ca^2+^ current due to its high Ca^2+^ permeability and prolonged open time of several hundred milliseconds (Kraft et al., 2004; Perraud et al., 2001; Sano et al., 2001). Nonetheless, more work directly linking TRPM2-Ca^2+^ influx in this pathway should be a priority for future investigation but remains technically challenging due to the lack of reliable antibodies directed at the channel (Cruz-Torres et al., 2020) and the poorly understood mechanism by which the channel is activated within the postsynaptic microdomain.

The Ca^2+^-activated signaling cascade downstream of TRPM2 ion channel activation is important to elucidate both for increased understanding of the mechanism of ischemia-induced alterations in E/I balance as well as future therapeutic development. Using two distinct pharmacological inhibitors of CaMKII, our data show that both Ca^2+^-stimulated and autonomous CaMKII activity are required for the elevation in postsynaptic GABAergic inhibition following GCI. Specifically, treatment with KN93, an inhibitor known to preferentially block Ca^2+^-dependent CaMKII (Vest et al., 2010), occludes the OGD effect on clustering (Fig 7F-G). In addition, our data also support the involvement of autonomous CaMKII in this mechanism, using the tatCN19o peptide which blocks both stimulated and autonomous CaMKII activity (Vest et al., 2010). CA/CPR induces elevated levels phosphorylated T286, and tatCN19o application reversed the CA/CPR-induced increase in sIPSC amplitude (Fig 6F_1_) and the OGD effect on postsynaptic GABAergic clustering (Fig 7C-D), further implicating the role of autonomous CaMKII activity, in addition to Ca^2+^-stimulated CaMKII activity, in our proposed pathway. These data are consistent with the literature implicating CaMKII in inhibitory synaptic potentiation. Autonomous CaMKII (T286phosphorylated) constitutively localizes to inhibitory synapses (Cook et al., 2022; Marsden et al., 2010) and intracellular application of CaMKII was shown to enhance GABAergic IPSCs (Wang et al., 1995; Wei et al., 2004). Additionally, CaMKII also undergoes autophosphorylation at T305 and T306 which is required for its translocation to inhibitory synapses within the dendritic shaft following NMDA-LTD stimulus (Cook et al., 2021). Following its movement to inhibitory synapses, CaMKII phosphorylates β2 or β3 subunits of GABA_A_Rs, resulting in GABA_A_R synaptic aggregation and inhibitory potentiation (Chiu et al., 2018; Petrini et al., 2014). Exploring whether CaMKII translocation to the inhibitory synapse and directly phosphorylates GABAergic proteins resulting in increased inhibitory function is an important avenue for future investigation.

Accumulating evidence suggests inhibitory LTP and homeostatic scaling up share downstream signaling pathways to regulate synaptic strength (Galanis and Vlachos, 2020; Vitureira and Goda, 2013). Exocytosis and subsequent lateral diffusion of GABA_A_Rs from the extrasynaptic membrane is a well-documented mechanism for rapidly fine-tuning inhibitory synaptic strength (Luscher et al., 2011). Multiple findings here support the notion that GABA_A_R clustering is increased through lateral diffusion. First, our data demonstrate that sIPSC amplitude and GABA_A_R clustering can be restored rapidly (within minutes to one hour). Second, we observe unaltered mRNA expression of GABA_A_R transcripts and protein levels of gephyrin post-CA/CPR, indicating no changes in the transcription and translation of GABAergic components at this timepoint (Fig S2). Nevertheless, translational upregulation of these components is necessary for the maintenance of inhibitory potentiation (Rajgor et al., 2020). We, therefore, cannot exclude the possibility that GCI alters these processes at more chronic timepoints. Lastly, our findings raise the intriguing possibility that increased inhibitory synaptic function represents a maladaptive homeostatic response to the acute loss of inhibition following excitotoxic insult. The OGD-reperfusion data hints at this possibility, revealing an initial acute reduction in the clustering of GABAergic components, followed by a steadily increasing and sustained increase in GABAergic synapses at later timepoints. The changes in the temporal dynamics of E/I balance between the acute and chronic phase warrant future investigation and may shed light into the relationship between early loss of GABAergic signaling and the delayed increase observed in our study.

In summary, this study uncovers a novel mechanism whereby enhanced GABAergic inhibition impairs excitatory synaptic plasticity in the context of cerebral ischemia, revealing the TRPM2-CaMKII pathway as a target for pharmacological intervention. This is particularly significant given the clinical challenges associated with direct GABA_A_R modulation. GABA_A_R antagonism induces epileptic seizures (Sperk et al., 2004), while GABA_A_R agonists largely failed in clinical trial as a treatment for acute ischemic stroke due to lack of efficacy and patients reporting numerous problematic side effects (Liu et al., 2018; Lyden et al., 2002; Wahlgren et al., 2000). Furthermore, our data shed light on the temporal changes in inhibitory synaptic function in response to ischemic insult. This observation may have profound impact on therapeutic window of some interventions, particularly GABA agonists, which have been shown to confer neuroprotection in animal models acutely (Liu et al., 2010; Liu et al., 2018) but, in light of this data, may be detrimental to functional recovery if administered at more chronic timepoints. Moreover, beyond ischemic injury, the pathway described here likely has broader implications considering that disruptions in synaptic accumulation of GABA_A_Rs are linked to numerous central nervous system disorders such as epilepsy, schizophrenia, depression, and substance abuse (Jacob et al., 2008; Mele et al., 2019). Further research into the precise molecular and cellular mechanisms underlying the influence of enhanced postsynaptic GABAergic function on excitatory LTP will be pivotal in developing targeted interventions to mitigate the long-term cognitive deficits associated with cerebral ischemia and various neurological diseases.

## ACKNOWLEDGEMENTS

This work was supported by an NIH Predoctoral NRSA F31NS120422 and T32GM763540 (A.B.), R01NS046072 (N.Q. and P.S.H.), R01NS118786 (P.S.H), R01NS092645 (P.S.H.), R01MH119154 (K.R.S.), R01MH128199 (K.R.S.), AHA Career Development Award (K.R.S.). J.D.G. was supported by grant T32GM763540, an AHA Predoctoral Fellowship (19PRE34380542) and an NIH Blueprint Diversity Specialized Predoctoral to Postdoctoral Advancement in Neuroscience (D-SPAN) Award (FNS120640A). We thank Ulrich Bayer for the gift of the tatCN19o peptide and Jacob Basak for critical reading of the manuscript. Graphical Abstract and OGD cartoons were created with BioRender.com.

## AUTHOR CONTRIBUTIONS

Conceptualization, A.M.B., H.O., and P.S.H.; Formal Analysis and Investigation, A.M.B., J.D.G., H.O., J.E.O., E.T., and N.C.; Methodology, E.T., and N.C.; Writing – Original Draft, A.M.B., H.O., N.Q., K.R.S., and P.S.H.; Writing – Review & Editing, A.M.B., N.Q., K.R.S., and P.S.H.; Supervision, N.Q., K.R.S., and P.S.H.; Funding Acquisition, A.M.B., J.D.G., N.Q., K.R.S., and P.S.H.

## DECLARATION OF INTERESTS

None.

## Notes

### Competing Interest Statement

The authors have declared no competing interest.

## REFERENCES

Alicke, B., Schwartz-Bloom, R. D., 1995. Rapid down-regulation of GABAA receptors in the gerbil hippocampus following transient cerebral ischemia. J Neurochem. 65, 2808–11.

Arancibia-Carcamo, I. L., et al., 2009. Ubiquitin-dependent lysosomal targeting of GABA(A) receptors regulates neuronal inhibition. Proc Natl Acad Sci U S A. 106, 17552–7.

Azad, T. D., et al., 2016. Neurorestoration after stroke. Neurosurg Focus. 40, E2.

Banks, M. I., et al., 1998. The synaptic basis of GABAA,slow. J Neurosci. 18, 1305–17.

Barcomb, K., et al., 2015. Live imaging of endogenous Ca(2)(+)/calmodulin-dependent protein kinase II in neurons reveals that ischemia-related aggregation does not require kinase activity. J Neurochem. 135, 666–73.

Cai, X., et al., 2023. TRPM2 regulates cell cycle through the Ca2+-CaM-CaMKII signaling pathway to promote HCC. Hepatol Commun. 7.

Carmichael, S. T., 2012. Brain excitability in stroke: the yin and yang of stroke progression. Arch Neurol. 69, 161–7.

Cheng, Y. D., et al., 2004. Neuroprotection for ischemic stroke: two decades of success and failure. NeuroRx. 1, 36–45.

Chiu, C. Q., et al., 2019. Preserving the balance: diverse forms of long-term GABAergic synaptic plasticity. Nat Rev Neurosci. 20, 272–281.

Chiu, C. Q., et al., 2018. Input-Specific NMDAR-Dependent Potentiation of Dendritic GABAergic Inhibition. Neuron. 97, 368–377 e3.

Clarkson, A. N., et al., 2010. Reducing excessive GABA-mediated tonic inhibition promotes functional recovery after stroke. Nature. 468, 305–9.

Cook, S. G., et al., 2021. CaMKII holoenzyme mechanisms that govern the LTP versus LTD decision. Sci Adv. 7.

Cook, S. G., et al., 2022. CaMKII T286 phosphorylation has distinct essential functions in three forms of long-term plasticity. J Biol Chem. 298, 102299.

Costa, A. C., Grybko, M. J., 2005. Deficits in hippocampal CA1 LTP induced by TBS but not HFS in the Ts65Dn mouse: a model of Down syndrome. Neurosci Lett. 382, 317–22.

Crosby, K. C., et al., 2019. Nanoscale Subsynaptic Domains Underlie the Organization of the Inhibitory Synapse. Cell Rep. 26, 3284–3297 e3.

Cruz-Torres, I., et al., 2020. Characterization and Optimization of the Novel Transient Receptor Potential Melastatin 2 Antagonist tatM2NX. Mol Pharmacol. 97, 102–111.

Deng, G., et al., 2017. Autonomous CaMKII Activity as a Drug Target for Histological and Functional Neuroprotection after Resuscitation from Cardiac Arrest. Cell Rep. 18, 1109–1117.

Dietz, R. M., et al., 2019. Reversal of Global Ischemia-Induced Cognitive Dysfunction by Delayed Inhibition of TRPM2 Ion Channels. Transl Stroke Res.

Dietz, R. M., et al., 2021. Functional Restoration following Global Cerebral Ischemia in Juvenile Mice following Inhibition of Transient Receptor Potential M2 (TRPM2) Ion Channels. Neural Plast. 2021, 8774663.

Escobar, I., et al., 2019. Altered Neural Networks in the Papez Circuit: Implications for Cognitive Dysfunction after Cerebral Ischemia. J Alzheimers Dis. 67, 425–446.

Galanis, C., Vlachos, A., 2020. Hebbian and Homeostatic Synaptic Plasticity-Do Alterations of One Reflect Enhancement of the Other? Front Cell Neurosci. 14, 50.

Garcia, J. D., et al., 2021. Stepwise disassembly of GABAergic synapses during pathogenic excitotoxicity. Cell Rep. 37, 110142.

Hill, K., et al., 2004. Inhibition of TRPM2 channels by the antifungal agents clotrimazole and econazole. Naunyn Schmiedebergs Arch Pharmacol. 370, 227–37.

Hiu, T., et al., 2016. Enhanced phasic GABA inhibition during the repair phase of stroke: a novel therapeutic target. Brain. 139, 468–80.

Jacob, T. C., et al., 2008. GABA(A) receptor trafficking and its role in the dynamic modulation of neuronal inhibition. Nat Rev Neurosci. 9, 331–43.

Jia, J., et al., 2011. Sex differences in neuroprotection provided by inhibition of TRPM2 channels following experimental stroke. J Cereb Blood Flow Metab. 31, 2160–8.

Joy, M. T., Carmichael, S. T., 2021. Encouraging an excitable brain state: mechanisms of brain repair in stroke. Nat Rev Neurosci. 22, 38–53.

Katz, A., et al., 2022. Pharmacologic neuroprotection in ischemic brain injury after cardiac arrest. Ann N Y Acad Sci. 1507, 49–59.

Kostandy, B. B., 2012. The role of glutamate in neuronal ischemic injury: the role of spark in fire. Neurol Sci. 33, 223–37.

Kraft, R., et al., 2004. Hydrogen peroxide and ADP-ribose induce TRPM2-mediated calcium influx and cation currents in microglia. Am J Physiol Cell Physiol. 286, C129–37.

Lake, E. M., et al., 2015. The effects of delayed reduction of tonic inhibition on ischemic lesion and sensorimotor function. J Cereb Blood Flow Metab. 35, 1601–9.

Lau, A., Tymianski, M., 2010. Glutamate receptors, neurotoxicity and neurodegeneration. Pflugers Arch. 460, 525–42.

Leao, R. N., et al., 2012. OLM interneurons differentially modulate CA3 and entorhinal inputs to hippocampal CA1 neurons. Nat Neurosci. 15, 1524–30.

Liu, B., et al., 2010. Preservation of GABAA receptor function by PTEN inhibition protects against neuronal death in ischemic stroke. Stroke. 41, 1018–26.

Liu, J., et al., 2018. Gamma aminobutyric acid (GABA) receptor agonists for acute stroke. Cochrane Database Syst Rev. 10, CD009622.

Luscher, B., et al., 2011. GABAA receptor trafficking-mediated plasticity of inhibitory synapses. Neuron. 70, 385–409.

Lyden, P., et al., 2002. Clomethiazole Acute Stroke Study in ischemic stroke (CLASS-I): final results. Stroke. 33, 122–8.

Marsden, K. C., et al., 2010. Selective translocation of Ca2+/calmodulin protein kinase IIalpha (CaMKIIalpha) to inhibitory synapses. Proc Natl Acad Sci U S A. 107, 20559–64.

Mele, M., et al., 2019. Alterations in GABA(A)-Receptor Trafficking and Synaptic Dysfunction in Brain Disorders. Front Cell Neurosci. 13, 77.

Mele, M., et al., 2014. GABA(A) receptor dephosphorylation followed by internalization is coupled to neuronal death in in vitro ischemia. Neurobiol Dis. 65, 220–32.

Moskowitz, M. A., et al., 2010. The science of stroke: mechanisms in search of treatments. Neuron. 67, 181–98.

Nakayama, S., et al., 2013. Sexually dimorphic response of TRPM2 inhibition following cardiac arrest-induced global cerebral ischemia in mice. J Mol Neurosci. 51, 92–8.

Neumann, J. T., et al., 2013. Global cerebral ischemia: synaptic and cognitive dysfunction. Curr Drug Targets. 14, 20–35.

Nusser, Z., et al., 1997. Differences in synaptic GABA(A) receptor number underlie variation in GABA mini amplitude. Neuron. 19, 697–709.

Orfila, J. E., et al., 2019. Delayed inhibition of tonic inhibition enhances functional recovery following experimental ischemic stroke. J Cereb Blood Flow Metab. 39, 1005–1014.

Orfila, J. E., et al., 2018. Cardiac Arrest Induces Ischemic Long-Term Potentiation of Hippocampal CA1 Neurons That Occludes Physiological Long-Term Potentiation. Neural Plast. 2018, 9275239.

Orfila, J. E., et al., 2014. Increasing small conductance Ca2+-activated potassium channel activity reverses ischemia-induced impairment of long-term potentiation. Eur J Neurosci. 40, 3179–88.

Perraud, A. L., et al., 2001. ADP-ribose gating of the calcium-permeable LTRPC2 channel revealed by Nudix motif homology. Nature. 411, 595–9.

Petito, C. K., et al., 1987. Delayed hippocampal damage in humans following cardiorespiratory arrest. Neurology. 37, 1281–6.

Petrini, E. M., et al., 2014. Synaptic recruitment of gephyrin regulates surface GABAA receptor dynamics for the expression of inhibitory LTP. Nat Commun. 5, 3921.

Rajgor, D., et al., 2020. Local miRNA-Dependent Translational Control of GABA(A)R Synthesis during Inhibitory Long-Term Potentiation. Cell Rep. 31, 107785.

Sano, Y., et al., 2001. Immunocyte Ca2+ influx system mediated by LTRPC2. Science. 293, 1327–30.

Schmidt-Kastner, R., Freund, T. F., 1991. Selective vulnerability of the hippocampus in brain ischemia. Neuroscience. 40, 599–636.

Schwartz-Bloom, R. D., Sah, R., 2001. gamma-Aminobutyric acid(A) neurotransmission and cerebral ischemia. J Neurochem. 77, 353–71.

Shimizu, T., et al., 2013. Androgen and PARP-1 regulation of TRPM2 channels after ischemic injury. J Cereb Blood Flow Metab. 33, 1549–55.

Smith, K. R., Kittler, J. T., 2010. The cell biology of synaptic inhibition in health and disease. Curr Opin Neurobiol. 20, 550–6.

Smith, K. R., et al., 2017. Differential regulation of the Rac1 GTPase-activating protein (GAP) BCR during oxygen/glucose deprivation in hippocampal and cortical neurons. J Biol Chem. 292, 20173–20183.

Sperk, G., et al., 2004. GABA and its receptors in epilepsy. Adv Exp Med Biol. 548, 92–103.

Steele, P. M., Mauk, M. D., 1999. Inhibitory control of LTP and LTD: stability of synapse strength. J Neurophysiol. 81, 1559–66.

Turski, L., et al., 1998. ZK200775: a phosphonate quinoxalinedione AMPA antagonist for neuroprotection in stroke and trauma. Proc Natl Acad Sci U S A. 95, 10960–5.

Udakis, M., et al., 2020. Interneuron-specific plasticity at parvalbumin and somatostatin inhibitory synapses onto CA1 pyramidal neurons shapes hippocampal output. Nat Commun. 11, 4395.

Verma, S., et al., 2012. TRPM2 channel activation following in vitro ischemia contributes to male hippocampal cell death. Neurosci Lett. 530, 41–6.

Vest, R. S., et al., 2010. Effective post-insult neuroprotection by a novel Ca(2+)/ calmodulin-dependent protein kinase II (CaMKII) inhibitor. J Biol Chem. 285, 20675–82.

Vitureira, N., Goda, Y., 2013. Cell biology in neuroscience: the interplay between Hebbian and homeostatic synaptic plasticity. J Cell Biol. 203, 175–86.

Vogels, T. P., et al., 2011. Inhibitory plasticity balances excitation and inhibition in sensory pathways and memory networks. Science. 334, 1569–73.

Wahlgren, N. G., Ahmed, N., 2004. Neuroprotection in cerebral ischaemia: facts and fancies--the need for new approaches. Cerebrovasc Dis. 17 Suppl 1, 153–66.

Wahlgren, N. G., et al., 2000. The clomethiazole acute stroke study (CLASS): Safety results in 1,356 patients with acute hemispheric stroke. J Stroke Cerebrovasc Dis. 9, 158–65.

Walters, M. R., et al., 2005. The AMPA antagonist ZK 200775 in patients with acute ischaemic stroke: a double-blind, multicentre, placebo-controlled safety and tolerability study. Cerebrovasc Dis. 20, 304–9.

Wang, Q., et al., 2017. Oxidative stress activates the TRPM2-Ca(2+)-CaMKII-ROS signaling loop to induce cell death in cancer cells. Biochim Biophys Acta Mol Cell Res. 1864, 957–967.

Wang, R. A., et al., 1995. Alpha-subunit of calcium/calmodulin-dependent protein kinase II enhances gamma-aminobutyric acid and inhibitory synaptic responses of rat neurons in vitro. J Neurophysiol. 73, 2099–106.

Wei, J., et al., 2004. Ca(2+)-calmodulin signalling pathway up-regulates GABA synaptic transmission through cytoskeleton-mediated mechanisms. Neuroscience. 127, 637–47.

Williams, L. E., Holtmaat, A., 2019. Higher-Order Thalamocortical Inputs Gate Synaptic Long-Term Potentiation via Disinhibition. Neuron. 101, 91–102 e4.

Wu, Q. J., Tymianski, M., 2018. Targeting NMDA receptors in stroke: new hope in neuroprotection. Mol Brain. 11, 15.

Xie, Y. F., et al., 2011. Dependence of NMDA/GSK-3beta mediated metaplasticity on TRPM2 channels at hippocampal CA3-CA1 synapses. Mol Brain. 4, 44.

