## Supplemental Information for "TRPM2-CaMKII signaling drives excessive GABAergic synaptic inhibition following ischemia"

**SUPPORTING INFORMATION**

| **Population Studies (different cells patched for drug conditions)** | | | | | | |
|---|---|---|---|---|---|---|
| **Internal Solution** | Condition | Mean Amplitude ±SEM | Mean Frequency ±SEM | Mean Decay ±SEM | N (mice) | n (cells) |
| **CsCl** | Sham | 50.55 ± 3.021 | 9.864 ± 0.6820 | 14.34 ± 1.328 | 8 | 24 |
| | Sham +tatM2NX | 48.63 ± 3.252 | 9.519 ± 1.109 | 13.36 ±1.528 | 12 | 19 |
| | CA/CPR | 70.75 ± 8.107 | 10.60 ± 0.9341 | 17.70 ± 1.955 | 11 | 17 |
| | CA/CPR +tatM2NX | 45.608 ± 2.207 | 8.553 ± 1.154 | 13.55 ± 1.550 | 10 | 18 |
| **Paired Studies (same cell before and after drug application)** | | | | | | |
| **Internal Solution** | Condition | Mean Amplitude ±SEM | Mean Frequency ±SEM | Mean Decay ±SEM | N (mice) | n (cells) |
| **CsCl** | Sham | 57.48 ± 4.498 | 8.389 ± 0.9211 | 16.66 ± 1.681 | 9 | 22 |
| | Sham +CTZ | 54.87 ± 4.923 | 8.298 ± 0.9358 | 17.73 ± 1.098 | | |
| | CA/CPR | 91.24 ± 8.188 | 9.504 ± 0.701 | 19.34 ± 1.073 | 11 | 21 |
| | CA/CPR +CTZ | 66.25 ± 6.011 | 8.061 ± 0.5825 | 20.74 ± 1.794 | | |
| **BAPTA** | Sham | 78.98 ± 10.29 | 7.507 ± 0.8206 | 18.96 ± 1.931 | 4 | 10 |
| | Sham +CTZ | 66.16 ± 7.021 | 6.677 ± 0.6352 | 15.46 ± 0.5528 | | |
| | CA/CPR | 66.45 ± 6.998 | 8.898 ± 0.8444 | 19.09 ± 1.387 | 7 | 10 |
| | CA/CPR +CTZ | 57.87 ± 7.862 | 8.131 ± 0.6292 | 18.94 ± 1.146 | | |
| **tatCN19o** | Sham | 67.29 ± 11.14 | 7.327 ± 1.021 | 22.88 ± 2.668 | 5 | 9 |
| | Sham +CTZ | 76.05 ± 11.2 | 8.143 ± 1.187 | 17.22 ± 1.909 | | |
| | CA/CPR | 65.56 ± 8.538 | 7.664 ± 0.8301 | 18.8 ± 1.838 | 4 | 9 |
| | CA/CPR +CTZ | 78.18 ± 19.51 | 7.73 ± 0.8476 | 18.45 ± 2.192 | | |

**Table S1. Summary of amplitude, frequency, tau decay for all conditions tested.**

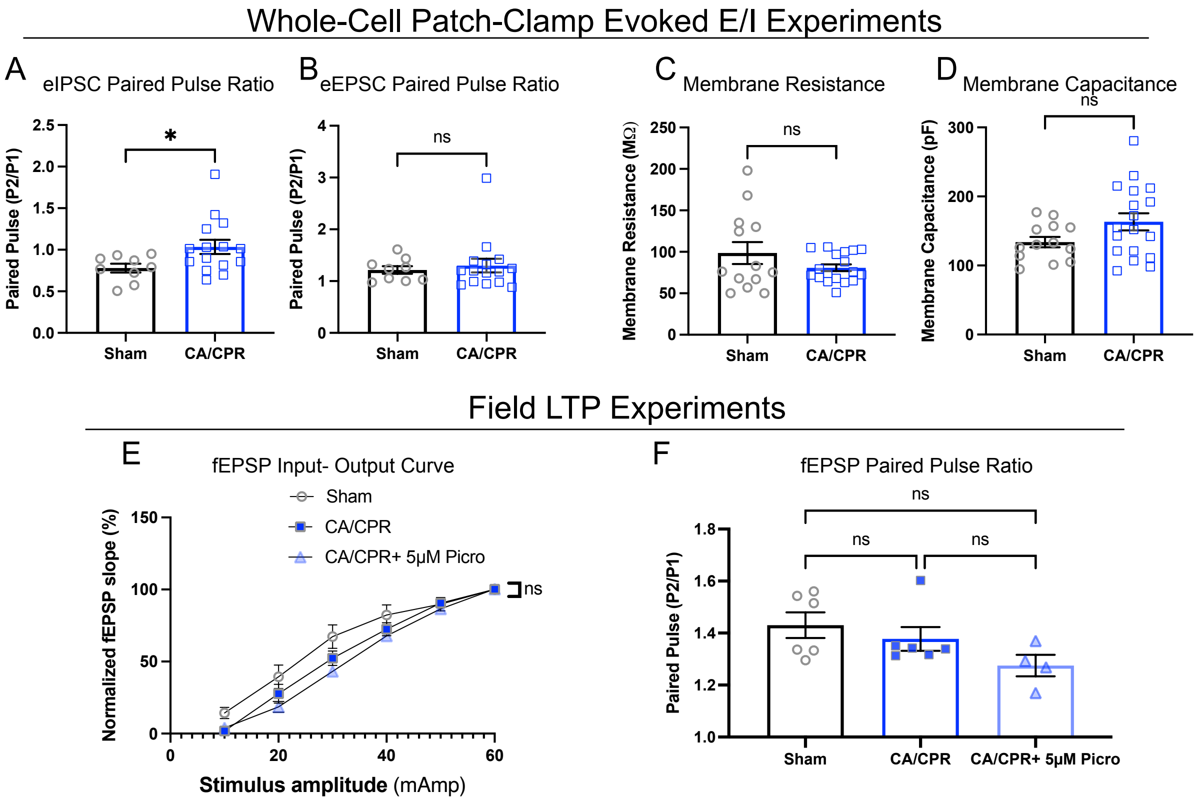


**Figure S1, related to Figure 1. Measurements of presynaptic function and membrane properties of CA1-pyramidal neurons (A-D) and CA1-Schaffer pathway (E-F).**

A. IPSC paired pulse ratio in CA1 pyramidal cells evoked at a holding potential of 0mV by inter-pulse interval of 100ms; n=11-15 cells/ 3-5 animals, unpaired t-test.

B. EPSC paired pulse ratio in CA1 pyramidal cells evoked at a holding potential of -70mV using inter-pulse interval of 50ms; n=11-15 cells/ 3-5 animals, unpaired t-test.

C. Membrane resistance of CA1 neurons measured immediately following cell break-in; n=13-18 cells/ 4-6 animals, unpaired t-test.

D. Membrane capacitance of CA1 neurons measured immediately following cell break-in; n=13-18 cells/ 4-6 animals, unpaired t-test

E. Input-output curve of fEPSP; n=4-5 slices/ 4-6 animals, simple linear regression, slope comparison.

F. Paired pulse ratio of fEPSP; n=4-5 slices/ 4-6 animals, One-Way ANOVA, Tukey's posthoc test.

Values represent mean ± SEM, *$p<0.05$.

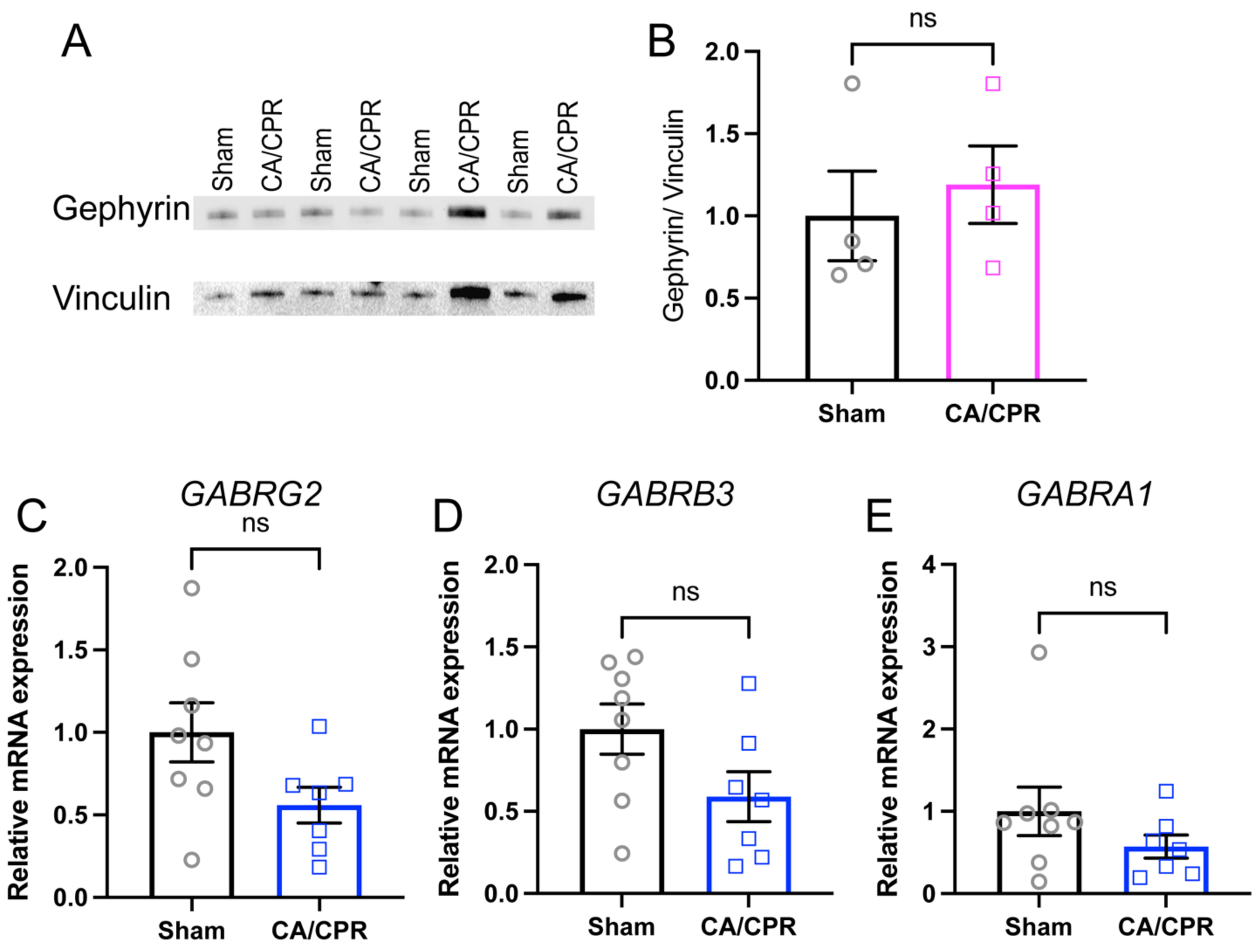


**Figure S2, related to Figure 3. Protein and mRNA expression of postsynaptic GABA components 7 days following CA/CPR.**

A. Western immunoblot measuring total levels of gephyrin normalized to βactin loading control in the P2 fraction from whole hippocampi.

B. Quantification of gephyrin levels normalized to mean sham.

C-E. Levels of mRNA transcript as measured by quantitative qPCR for genes encoding the $GABA_AR$-γ2 (C), -β3 (D), -α1 (E) subunits.

Values represent mean ± SEM.

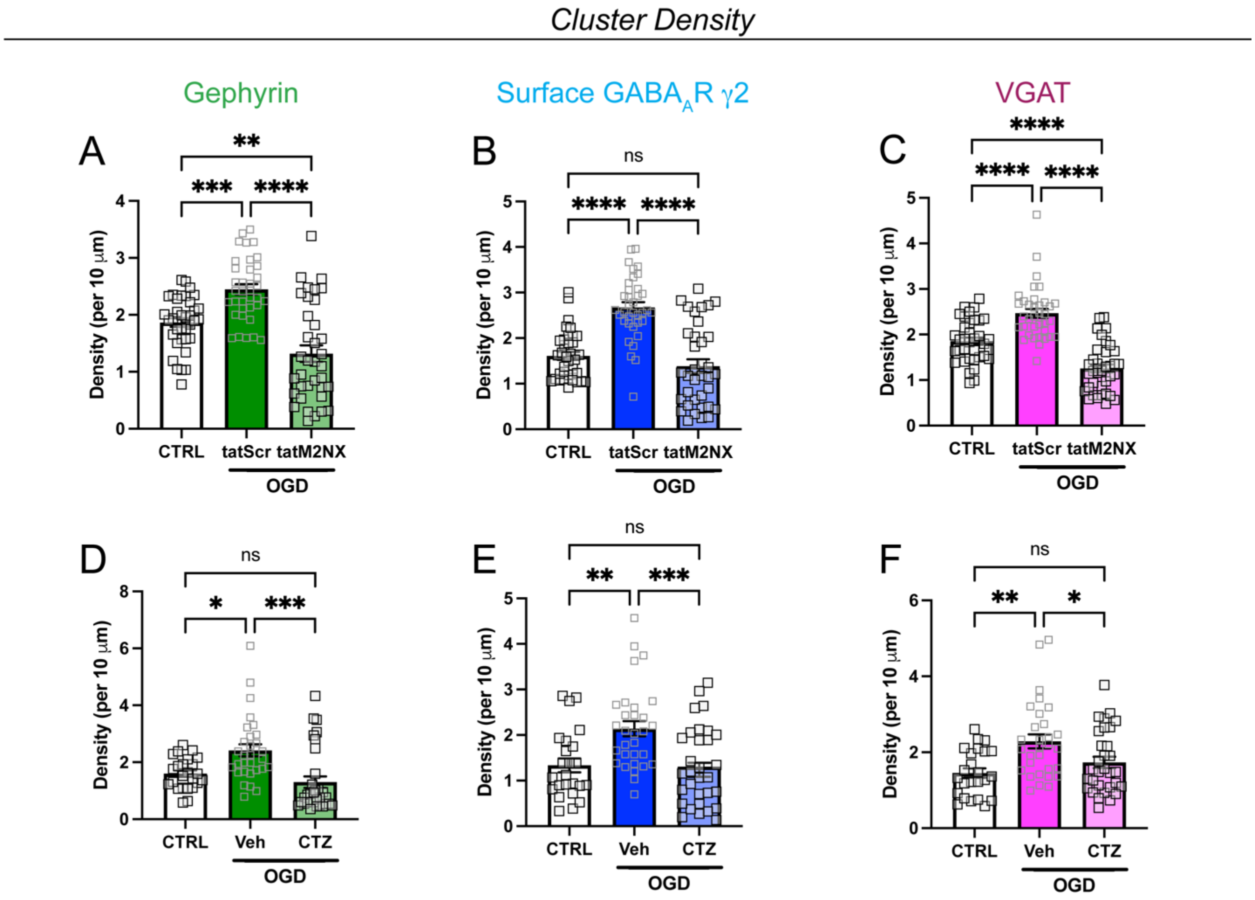


**Figure S3, related to Figure 5. TRPM2 ion channel inhibition reduceS cluster density of synaptic GABA proteins following OGD.**

A-C. Quantification of cluster density following treatment with tatM2NX for (A) gephyrin, (B) surface GABA$_A$R-γ2, and (C) VGAT; n=30-36 neurons per condition; One-Way ANOVA, Tukey's posthoc.

D-F. Quantification of cluster density following treatment with CTZ for (A) gephyrin, (B) surface GABA$_A$R-γ2, and (C) VGAT; n=30-36 neurons per condition; One-Way ANOVA, Tukey's posthoc.

Values represent mean ± SEM. *p<0.05; ** p<0.01, ***p<0.001, ****p<0.0001.

*Cluster Density*

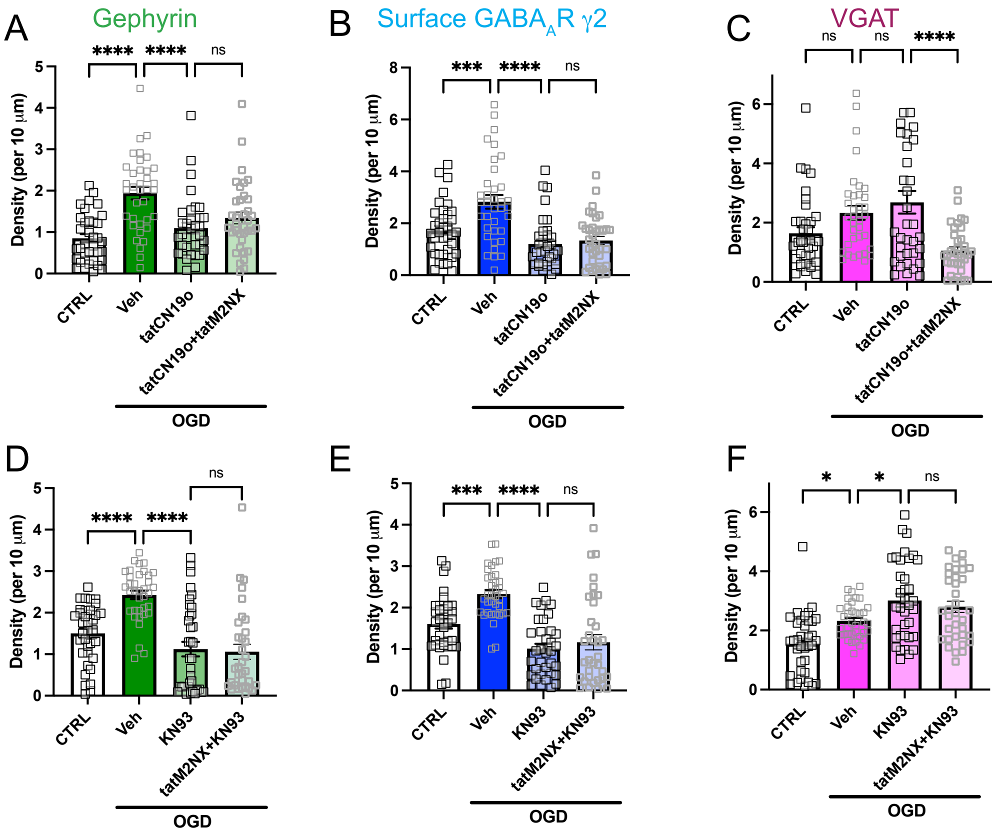


**Figure S4, related to Figure 7. Inhibition of $Ca^{2+}$-dependent CaMKII activity reduces cluster density of synaptic GABA components following OGD.**

A-C. Quantification of cluster density following treatment with tatCN19o or tatCN19o+ tatM2NX for (A) gephyrin, (B) surface GABA$_A$R-γ2, and (C) VGAT; n=30-36 neurons per condition; One-Way ANOVA, Tukey's posthoc.

D-F. Quantification of cluster area following treatment with KN93 or KN93+tatM2NX for (D) gephyrin, (E) surface GABA$_A$R-γ2, and (F) VGAT; n=30-36 neurons per condition; One-Way ANOVA, Tukey's posthoc.

Values represent mean ± SEM. *p<0.05; ** p<0.01, ***p<0.001, ****p<0.0001.

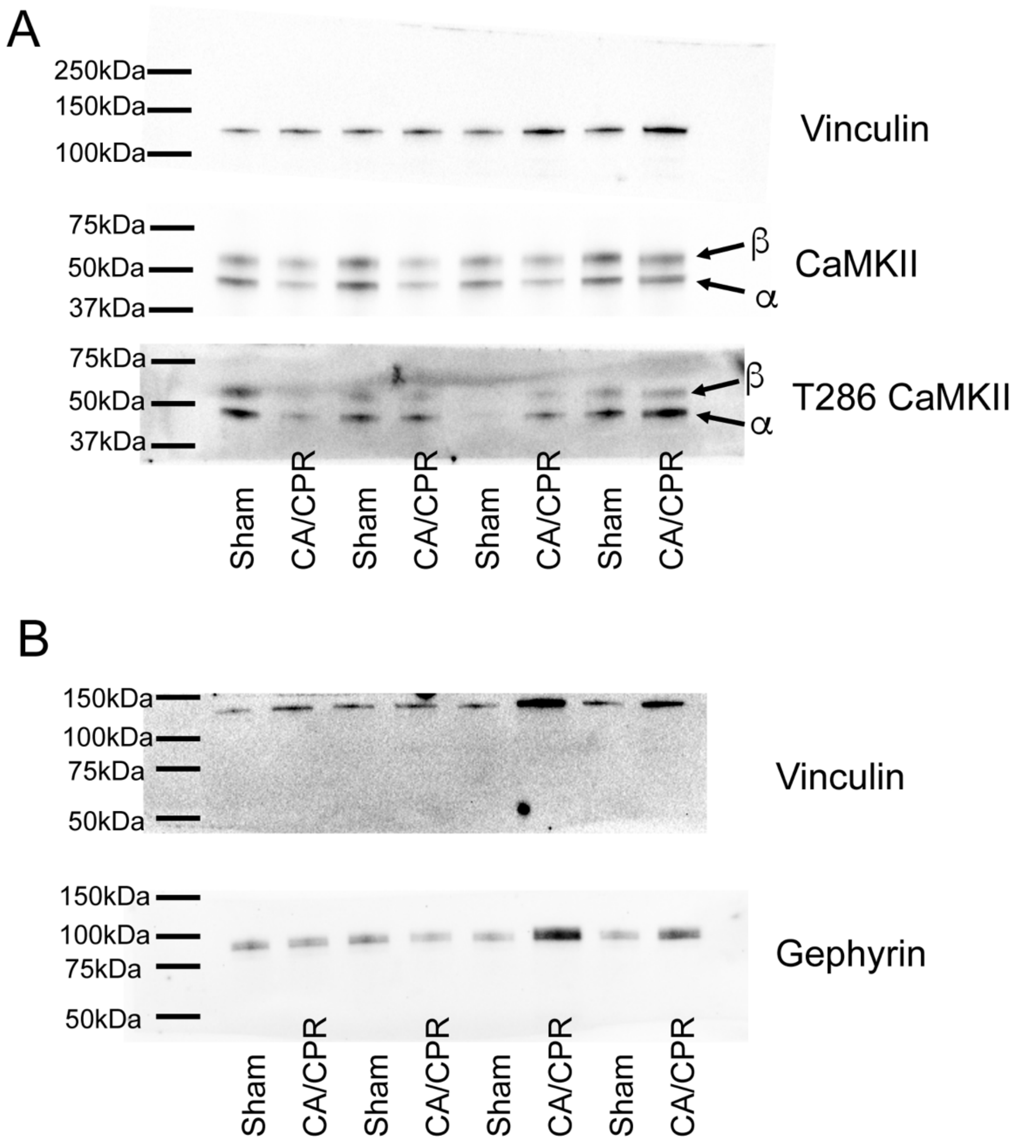


**Figure S5, related to Figure 6 and Figure S2. Raw images of Western immunoblots with corresponding molecular weight ladders.**

A. Images of CaMKII blot from Figure 6. T286 CaMKII blot is the stripped total CaMKII blot.

B. Images of gephyrin blot from Figure S2.
